# The JNK and Hippo pathways control epithelial integrity and prevent tumour initiation by regulating an overlapping transcriptome

**DOI:** 10.1101/2024.06.19.599466

**Authors:** Katrina A. Mitchell, Joseph H. A. Vissers, Jonathan M. Pojer, Elliot Brooks, Abdul Jabbar Saiful Hilmi, Anthony T. Papenfuss, Jan Schröder, Kieran F. Harvey

## Abstract

Epithelial organs maintain their integrity and prevent tumour initiation by actively removing defective cells, such as those that have lost apicobasal polarity. Here, we identify how transcription factors of two key signalling pathways – Jun-N-terminal kinase (JNK) and Hippo – regulate epithelial integrity by controlling transcription of an overlapping set of target genes. Targeted DamID experiments reveal that in proliferating cells of the *Drosophila melanogaster* eye, the AP-1 transcription factor Jun, and the Hippo pathway transcription regulators Yorkie and Scalloped bind to a common suite of target genes that regulate organ growth. In defective neoplastic cells, AP-1 transcription factors repress transcription of growth genes together with the CtBP co-repressor. If gene repression by AP-1/CtBP fails, neoplastic tumour growth ensues, driven by Yorkie/Scalloped. Thus, AP-1/CtBP eliminates defective cells and prevents tumour initiation by acting in parallel to Yorkie/Scalloped to repress expression of a shared transcriptome. These findings shed new light on the maintenance of epithelial integrity and tumour suppression.

## INTRODUCTION

The most common cell type in animal organs are epithelial cells, which are essential for the barrier function of external and internal organs. Epithelial cells are polarised and form specialised apical, basal and lateral surfaces that mediate cellular adhesion and organ architecture ^1^. During both development and homeostasis, damaged and defective cells are actively removed from epithelia in order to maintain organ integrity and prevent tumour initiation ^2^. In *Drosophila melanogaster* epithelial tissues, aberrant cells, such as those with defective apicobasal polarity, are eliminated following induction of the TNF pathway (also known as the JNK pathway) ^3^. JNK pathway activity is controlled by extracellular ligands and transmembrane receptor proteins; in *D. melanogaster* the TNF ligand is named Eiger and the receptors are Grindelwald and Wengen ^3^. The JNK pathway controls transcription by regulating the activity of the heterodimeric AP-1 transcription factor pair Jun and Fos. In humans, there are multiple Jun and Fos family members, whilst in *D. melanogaster* AP-1 is comprised of a sole Jun protein (also known as Jun-related antigen – Jra) and a sole Fos protein (also known as Kayak – Kay). AP-1 transcription factors can also be regulated by the Epidermal growth factor (EGFR) pathway ^3–5^. Elimination of polarity-defective cells from developing epithelia has been reported to be induced by cell competition signals from neighboring cells ^6^ and, more recently, by cell-autonomous TNF/JNK signalling ^7^. If TNF/JNK pathway activity is perturbed (e.g., by mutating the gene encoding the TNF ligand *eiger*, or by expressing a dominant negative version of the Jun N-terminal kinase Basket), polarity-defective tissue not only survives but can grow overtly and form tumours ^8–11^.

JNK signalling has been reported to control epithelial integrity in concert with other signalling pathways, including Hippo, which a key regulator of organ growth. The Hippo pathway responds to mechanical, cell-cell adhesion and cell polarity cues, which are conveyed by several upstream pathway proteins ^12–15^. These converge on and activate a core kinase cassette [in *D. melanogaster*: Hippo (Hpo), Salvador (Sav), Mats and Warts (Wts)] ^16–25^. Wts is the most downstream kinase in the Hippo pathway and regulates transcription by phosphorylating the Yorkie (Yki) transcriptional co-activator protein ^26,27^, and controlling the rate of Yki nucleo-cytoplasmic shuttling ^28^. Yki regulates transcription by partnering with the Scalloped (Sd) DNA binding protein ^29–32^.

Several studies have reported that the JNK pathway directly activates the Hippo pathway to suppress tissue growth. This has been concluded in the context of cell polarity loss as well as overexpression of JNK pathway genes, both of which induce JNK pathway activity. The mechanism by which JNK and Hippo co-regulate tissue growth is controversial however because in different experimental scenarios JNK has been reported to both activate and repress Hippo pathway activity. Specifically, in cells where the cell polarity protein Lgl was depleted or the JNK pathway was activated by expressing a hyperactive *hep* transgene, the JNK pathway was reported to block Hippo pathway activity by phosphorylating Ajuba (Jub) and enhancing its ability to bind and sequester Wts, and thereby promote Yki activity ^33,34^. By contrast, stimulation of JNK signalling by Eiger overexpression was reported to induce Wts activity and thereby limit Yki/Sd-dependent transcription and hence tissue growth ^35^.

In contrast to these pathway crosstalk studies the Hippo and JNK pathway transcription factors have been found to act in parallel to co-regulate transcription in numerous settings. Unbiased transcription factor binding studies of both YAP and TEAD (the mammalian Yki and Sd orthologues, respectively) in human cells identified AP-1 binding motifs as being highly enriched in genomic regions that were bound by YAP and TEAD ^36–38^. Similar observations were reported in *D. melanogaster* epithelial tissues, i.e., enrichment of the AP-1 motif in the putative regulatory regions of Sd target genes ^39^. Functional studies in cultured cells, organoids and tumour xenografts also revealed that AP-1 and YAP/TEAD regulate a subset of common target genes and cooperatively promote the ability of cells to grow in soft agar and as xenografted tumours in mice ^36,40^, and cell migration and invasion ^38,41^. In mice, AP-1 and TEAD4 also cooperate to determine whether cells adopt the hemogenic or vascular smooth muscle fates ^42^. In *D. melanogaster,* AP-1 and Yki/Sd cooperatively control the differentiation of alary muscle cells during development ^43^, while in epithelial tissues with defective polarity, Yki/Sd and the AP-1 transcription factor Fos co-regulate transcription of the *unpaired3 (upd3)* gene ^44^.

To further interrogate the mechanism by which the JNK and Hippo pathways regulate epithelial tissue growth, we generated new *D. melanogaster* loss of function alleles of genes that encode for the AP-1 transcription factor proteins, Jun and Fos, and surveyed the genome binding regions of Jun, Yki and Sd, in proliferating eye cells using targeted DamID. Our data indicates that the predominant mode by which the JNK and Hippo pathways co-regulate neoplastic tissue growth is by controlling the transcription of common target genes, rather than by pathway crosstalk at the level of regulatory kinases that act upstream of Yki/Sd and AP-1. Furthermore, we find no evidence that the JNK pathway transcription factors regulate the growth of eye imaginal discs that are growing normally or that overproliferate because of Yki hyperactivation. In clonal neoplasia, JNK signalling limits clone growth by repressing transcription of shared Yki/Sd/AP-1 target genes via AP-1 and the CtBP co-repressor. Thus, JNK acts in parallel to Hippo to repress expression of a shared transcriptome to maintain epithelial integrity and prevent tumour initiation.

## RESULTS

### JNK pathway transcription factors limit neoplastic tumour growth in parallel to the Hippo pathway

Multiple studies have reported that to maintain epithelial integrity, the JNK pathway is activated in aberrant cells and triggers their removal by activating the Hippo pathway. Increased Hippo activity then limits cell proliferation and viability by repressing Yki and its ability to promote transcription ^9,35^. Given that the predominant mode by which signalling pathways modulate transcription and hence cell behaviour is via their canonical transcription factors, rather than via crosstalk with other signalling pathways, we re-examined these claims. A tissue growth setting where the TNF/JNK pathway is known to have a robust role is in the context of neoplasia resulting from the loss of apicobasal polarity regulators such as Scribble (Scrib)^3^. In *scrib* imaginal disc clones, JNK pathway activity is robustly induced and represses clone growth ^8^, whilst blockade of upstream TNF/JNK signalling dramatically reverses this phenotype and allows *scrib* clones to substantially overgrow ^8–11^. Several studies concluded that JNK limits the growth of *scrib* clones by repressing Yki activity and access to the nucleus ^9,35^, rather than by acting through its canonical transcription factors Jun and Fos, while one study showed that RNAi-mediated depletion of Fos can suppress apoptosis in *scrib* clones ^10^. An independent study also reported that Yki was more nuclear and active in epithelial tissues with disrupted apicobasal polarity^33^. Given these conflicting observations, we examined this further, by using CRISPR/Cas9 genome editing to generate new mutant alleles of both *jun* and *fos*. These new alleles were embryonic lethal and exhibited dorsal closure phenotypes similar to published *jun* and *fos* alleles, indicating that they are strong loss of function alleles and likely to be null alleles (Figures S1A-S1C).

The JNK pathway’s role in imaginal disc growth is controversial; early studies of tissue lacking the *basket* gene (which encodes the JNK protein) found no obvious impact on eye growth ^45^, while, more recently, the JNK pathway was reported to control wing growth, but to do so independent of AP-1, by modulating Yki activity ^46^. Using our new *jun* and *fos* alleles, we found that eye tissue harbouring mutations in either, or both, *jun^KM^* and *fos^KM^,* grew normally (Figures S1D-S1F). To investigate a role for *jun* in the growth of *scrib* clones, we generated larval eye imaginal discs that harboured clones for the right arms of both chromosomes 2 and 3. In this scenario, a complex set of tissues of different genotypes is generated: tissue that is homozygous mutant for both *jun^KM^* and *scrib*^1^ lacks both RFP and GFP, homozygous *jun^KM^* tissue lacks GFP, homozygous *scrib*^1^ tissue lacks RFP and tissue that is heterozygous for one or both of these genes or homozygous wild-type will express either RFP or GFP, or both. Loss of *scrib* reduced clone growth, while, as above, *jun^KM^* clones grew normally (Figures 1A, 1B and 1D). Loss of *jun* partially restored the growth of *scrib*^1^ clones suggesting that it restricts the growth potential of eye tissue upon JNK pathway induction (Figures 1C-1D). To examine this further, we tested whether Fos is required for the JNK pathway to limit the growth of *scrib*^1^ clones, by generating animals with a recombinant chromosome harbouring FRT82B and both *scrib*^1^ and *fos^KM^* alleles. While *scrib*^1^ eye clones undergrew, *scrib*^1^*, fos^KM^* double mutant eye clones were not only rescued, but dramatically overgrew (Figures 1E-1I). This was observed for two independent *scrib*^1^*, fos^KM^* recombinant strains, as well as a strain generated with a published hypomorphic *fos* allele (*scrib*^1^*, fos*^2^) (Figure S1G). Note, the restoration of *scrib*^1^ clone growth by *fos^KM^*, was more striking than for *jun^KM^*, because more *scrib*^1^*, fos^KM^* double mutant tissue is generated in this simpler genetic experiment than in the *scrib*^1^*, jun^KM^* experiment. Together, these experiments indicate that the AP-1 transcription factors Jun and Fos limit the growth of neoplastic *scrib* cells.

**Figure 1.**
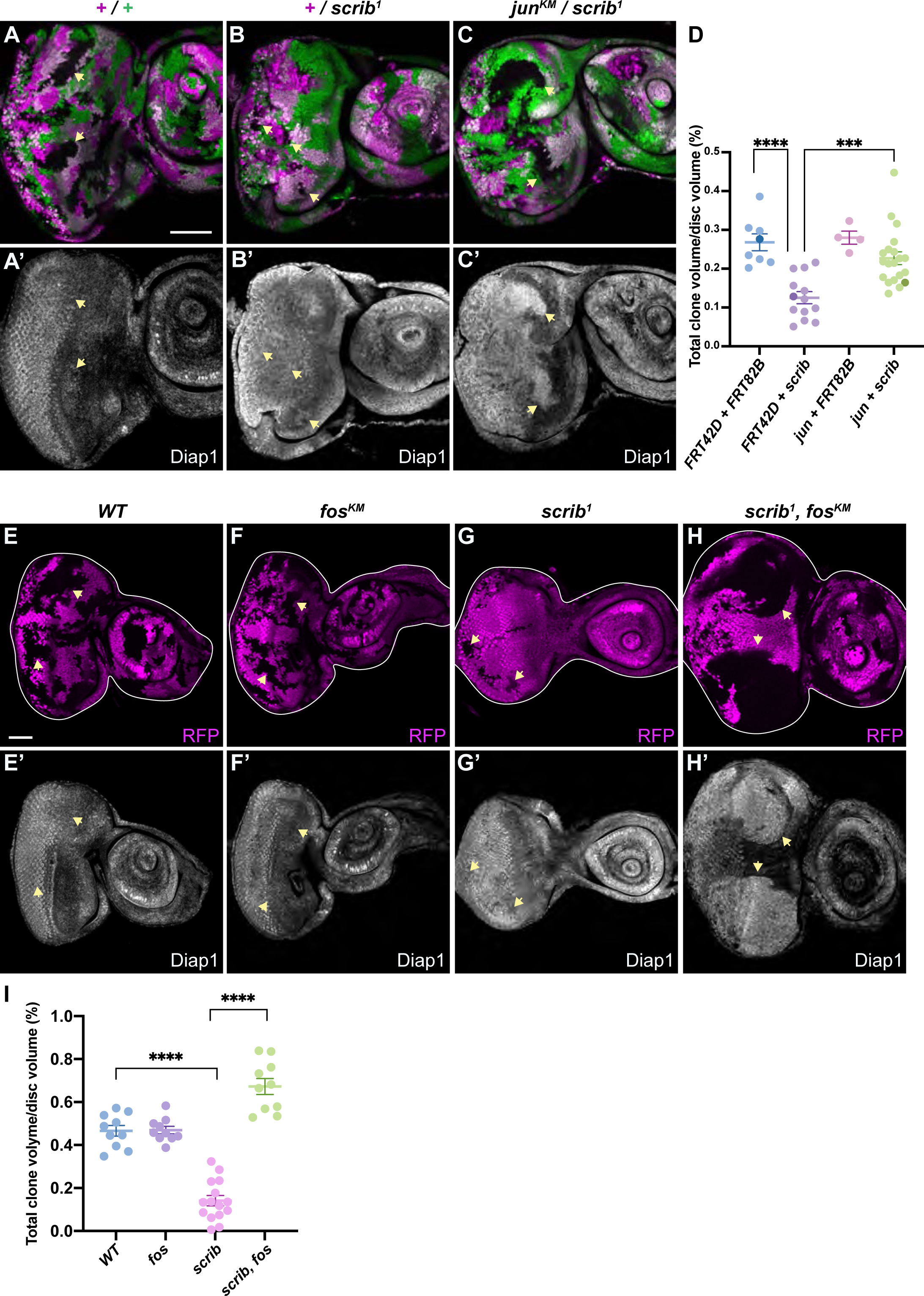
Fos and Jun limit neoplastic clone growth and DIAP1 expression. **A-C’.** Mosaic third instar larval eye-antennal discs containing control tissue marked by RFP (FRT82B-derived tissue, magenta) and/or GFP (FRT42D-derived tissue, green), or tissue lacking both markers - wild-type in (A), *scrib*^1^ in (B) and *jun^KM^* and *scrib*^1^ in (C). The bottom row of panels (A’-C’) show DIAP1 expression (grayscale) in the same tissues. Arrowheads indicate selected clones, the scale bar represents 50μm. **D.** Chart showing quantification of clone sizes of the genotypes in (A-C), shown as a ratio of RFP^-^ volume or GFP^-^ volume, over total eye-antennal disc volume. The overlapping clone genotypes are: *FRT42D wildtype and FRT82B wildtype*; *FRT42D wildtype and FRT82B scrib*^1^; *FRT42D jun^KM^* and *FRT82B wildtype*; and *FRT42D jun^KM^* and *FRT82B fos^KM^*. n = 8, 13, 4, 22, respectively. Data are represented as mean ± SEM. p-values were obtained using unpaired t-tests. *** p < 0.001, **** p < 0.0001. **E-H’.** Mosaic third instar larval eye-antennal discs containing clones marked by the absence of RFP (magenta) of the following genotypes: *wild-type* (E), *fos^KM^* (F), *scrib*^1^ (G), or *scrib*^1^*, fos^KM^* (H). The bottom row of panels (E’-H’) show DIAP1 expression (grayscale) in the same tissues. Arrowheads indicate selected clones, the scale bar represents 50μm. **I.** Chart showing quantification of clone sizes of the genotypes in (E-H), over total eye-antennal disc volume. n = 10, 10, 15, 10, respectively. Data are represented as mean ± SEM. p-values were obtained using unpaired t-tests. **** p < 0.0001.

These findings argue that JNK pathway activation limits neoplastic growth via its canonical transcription factors, i.e., AP-1, rather than by activating the Hippo pathway to repress Yki/Sd-regulated transcription. To test this further, we assessed the expression of Hippo pathway target genes, as well as localization of Yki, which accumulates in the nucleus if Hippo signalling is defective ^26,27^. If the JNK pathway indeed stimulates the Hippo pathway in neoplastic cells, then Yki/Sd target genes should be repressed in both *scrib* and *scrib, fos* clones and Yki should be more cytoplasmic (i.e., inactive). However, we observed results that directly opposed these predictions. The protein encoded by the Yki/Sd target gene *Diap1* was in fact elevated in both *scrib*^1^, *jun^KM^* tissue and *scrib*^1^, *fos* clones, rather than repressed (Figures 1A’-1C’, 1E’-1H’ and S1G). This result was confirmed in *scrib*^1^, *fos^KM^* clones using an enhancer trap line (*Diap1-LacZ*) that reports activity of the *Diap1* gene (Figures 2A-2C’ and 2M). In addition, neither DIAP1, nor *Diap1-lacZ* were altered in *scrib*^1^ or *fos^KM^* single mutant clones (Figures 1E-1G, 1J, 2A-2B’ and 2M). An independent Yki/Sd target gene *expanded* (*ex*, assessed with *ex-LacZ,* a transcription reporter), was also elevated in *scrib*^1^, *fos^KM^* clones in both eye-antennal and wing discs, but not the single mutant clones (Figures 2D-2F’, 2N and S1H). Finally, we assessed two transcription reporters of the *bantam* (*ban*) gene, *ban3-GFP* and *brC12-lacZ*, that are based on small enhancer fragments (1.2kb and 0.4kb) upstream of *ban* and are regulated by both Yki and Sd ^47–49^. Both *ban3-GFP* and *brC12-lacZ* were elevated in *scrib*^1^, *fos^KM^* clones but were unchanged in *scrib*^1^ clones and *fos^KM^* clones (Figures 2G-2L’, 2O and 2P).

**Figure 2.**
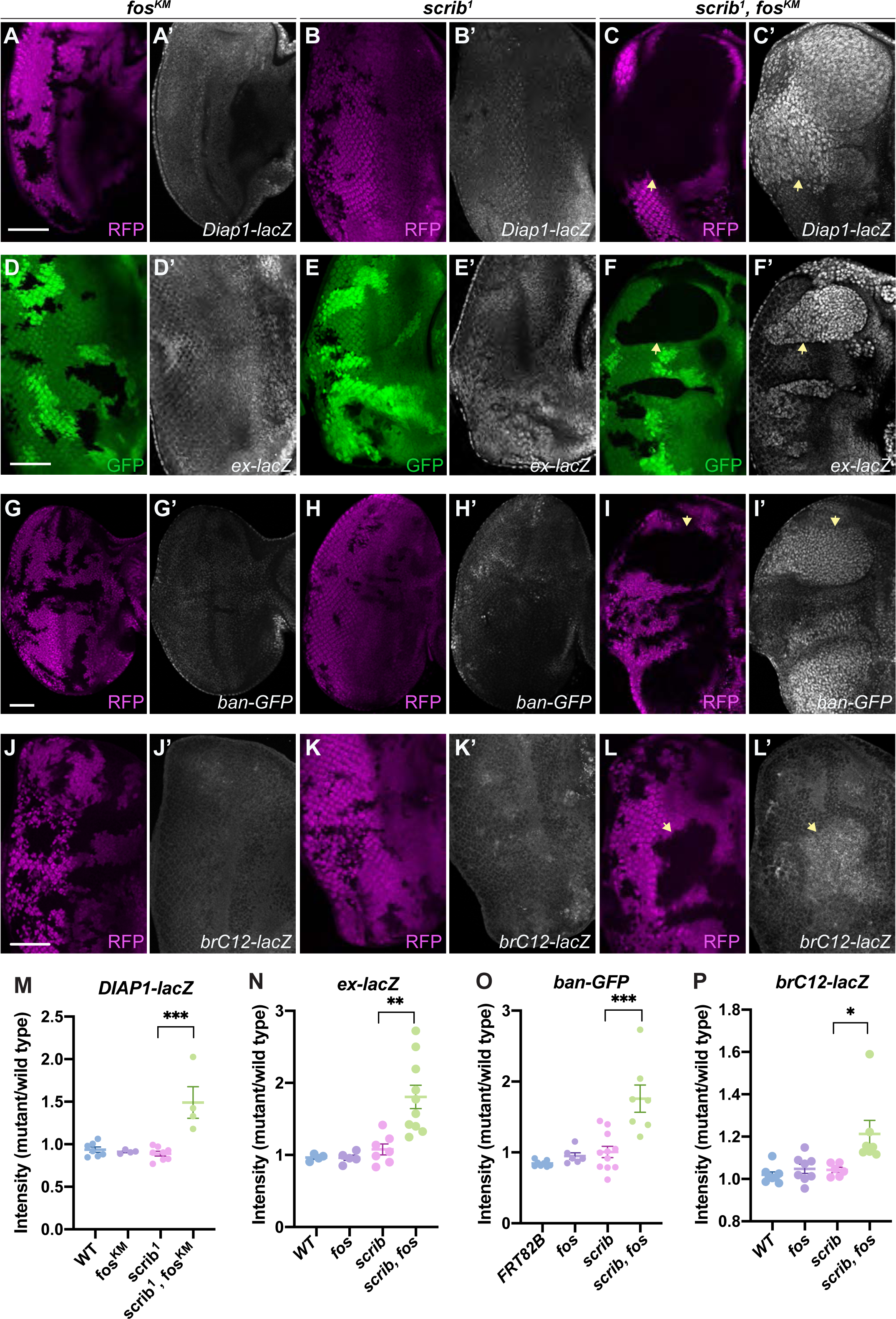
Fos limits expression of Hippo pathway target genes in neoplastic cells. **A-L’.** Mosaic third instar larval eye-antennal discs containing clones marked by the absence of RFP (magenta) or GFP (green) of the following genotypes: *fos^KM^* (E, D, G, J), *scrib*^1^ (B, E, H, K), or *scrib*^1^*, fos^KM^* (C, E, H, K). Tissues were stained with anti-β-Gal (greyscale) to reveal *Diap1-lacZ* (A’-C’), *ex-LacZ* (D’-F’) or *brC12-LacZ* (J’-L’). *ban-GFP* (greyscale) is in G’-I’. Arrows indicate selected clones, scale bars represent 50μm. **M-P.** Charts showing quantification of *Diap1-lacZ* (M), *ex-lacZ* (N), *ban-GFP* (O) or *brC12-lacZ* (P) in mutant versus wild-type larval eye disc tissue of the indicated genotypes. n = 4, 4, 8, 4 in (M), 4, 5, 7, 10 in (N), 8, 6, 11, 7 in (O), and 8, 8, 6, 7 in (P). Data are represented as mean ± SEM. p-values were obtained using unpaired t-tests. * p < 0.05, ** p < 0.01, *** p < 0.001.

The finding that expression of multiple Yki/Sd target genes were elevated, as opposed to repressed, in *scrib*^1^, *fos^KM^* clones argues that Yki is not in fact repressed by the JNK pathway in neoplastic cells, because the proposed JNK-mediated repression of Yki should still be intact in these cells. It also suggests that Fos somehow hinders Yki activity in *scrib* cells, because in the absence of Fos, Yki/Sd target genes were elevated. Consistent with this, Yki was more nuclear in both *scrib*^1^ and *scrib*^1^*, fos^KM^* clones, than in neighbouring control tissues and Yki abundance was also elevated (Figures 3A-3F). Yki nuclear abundance was also elevated in eye tissue harbouring an independent *scrib* allele (*scrib*^2^) (Figure S1I). These findings are striking because the growth properties of *scrib*^1^ and *scrib*^1^*, fos^KM^* clones are dramatically different, i.e., *scrib*^1^ clones undergrow and eventually die, whilst *scrib*^1^*, fos^KM^* clones overgrow considerably (Figures 1G-1I). This further supports the hypothesis that the key mediator of JNK’s ability to suppress neoplastic clone growth is AP-1-regulated transcription, rather than Hippo pathway activation and subsequent Yki inhibition. Furthermore, this data indicates that the Hippo activity pathway is compromised in *scrib* clones, rather than activated.

**Figure 3.**
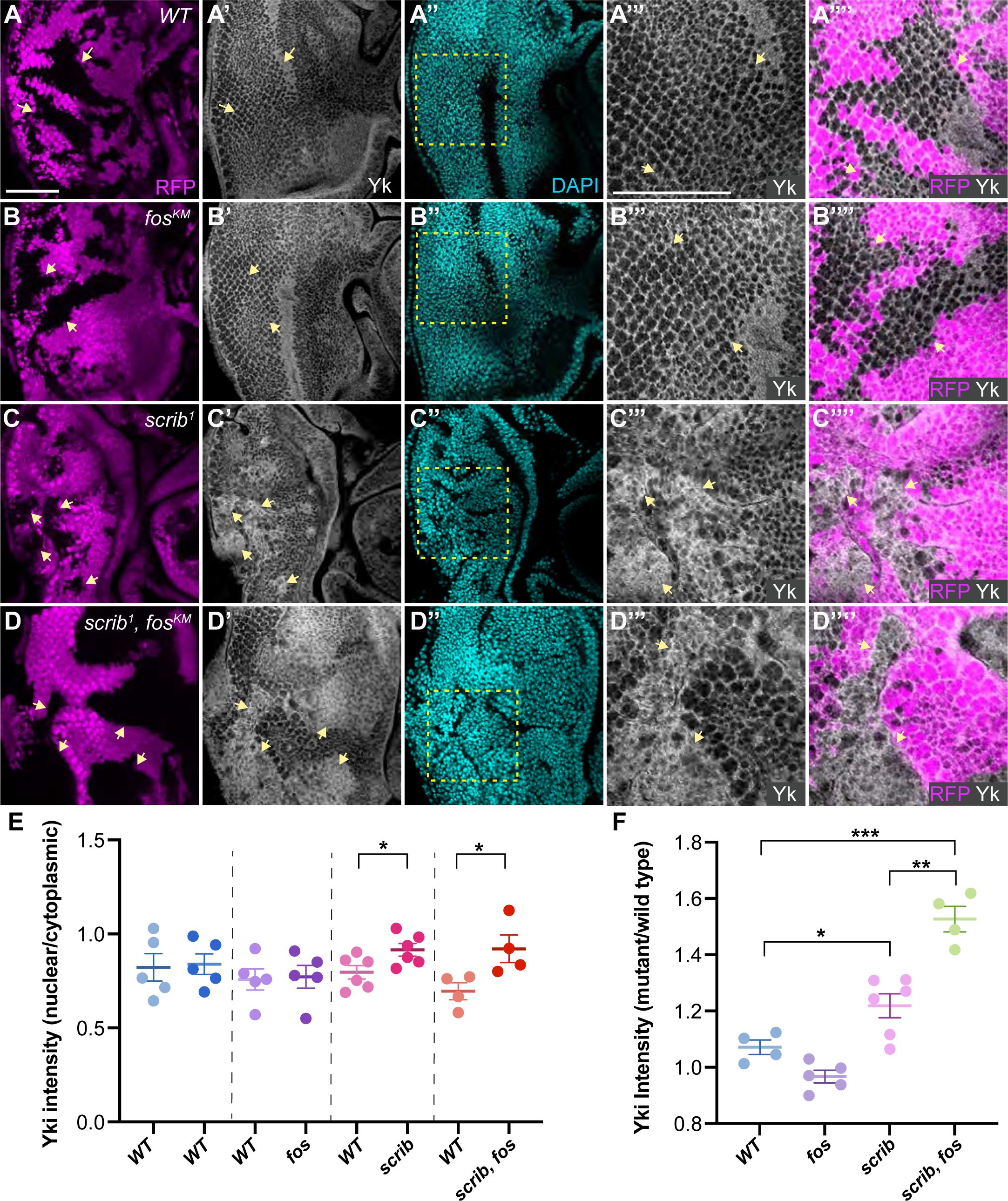
Epithelial cells with defective apicobasal polarity have impaired Hippo pathway activity. **A-D.** Mosaic third instar larval eye-antennal discs stained with anti-Yki (greyscale). RFP (magenta) marks control wild-type tissue while non-RFP tissue is either wild-type (A) or *fos^KM^* (B), *scrib*^1^ (C), or *scrib*^1^*, fos^KM^* (D). DAPI is cyan, arrows indicate select clones and the boxed region in A”-D” is shown at higher magnification in A’’’-D’’’ and the merged images in A’’’’-D’’’’. Scale bars represent 50μm. **E-F.** Charts showing the ratio of Yki intensity in the nucleus versus the cytoplasm (E), and in mutant tissue of the indicated genotypes versus wildtype tissue (F), in larval eye discs. n = 5, 5, 5, 5, 6, 5, 4, 4 in (E), and 4, 5, 6, 4 in (F). Data are represented as mean ± SEM. p-values were obtained using unpaired t-tests. * p < 0.05, ** p < 0.01, *** p < 0.001.

### Hippo and JNK pathway transcription factors share many target genes in proliferating eye cells

The results described above indicate that JNK induction limits epithelial tissue growth via the AP-1 transcription factors. In addition, it shows that JNK induction in *scrib* clones dominates the growth-promoting potential of increased nuclear Yki. To investigate the mechanism underlying these observations, we sought to identify the target genes of AP-1 and Yki/Sd in the growing eye. For this, we employed targeted DamID (TaDa), which allows in vivo profiling of genome binding by transcription factors, with precise spatiotemporal control ^50^. *Yki-Dam, Dam-Sd, Jun-Dam* or the *Dam* control transgene were expressed in developing *D. melanogaster* eyes for 24 hours during a developmental period where undifferentiated progenitor cells actively proliferate ^51^. TaDa revealed Yki binding to 2,705 genes and Sd to 2,9690 genes, with the majority of these being bound by both proteins (Figures 4A and S2A-S2B and Table S1). The Yki TaDa dataset overlapped significantly with a Yki ChIP-seq dataset from late third instar larval eye discs ^52^, while the Sd TaDa dataset overlapped significantly with Sd ChIP-seq datasets from both eye and wing discs ^52,53^ (Figure S2C). To determine which transcription factor motifs were enriched in the Yki, Sd and Jun TaDa experiments, we employed the commonly used Hypergeometric Optimization of Motif EnRichment (HOMER) analysis ^54^. This revealed that the most significantly enriched DNA binding motifs within shared Yki/Sd genome binding regions were those of Grainyhead (Grh), TEAD-Sd and AP-1 (Figure 4A). Jun bound to 3,460 genes, with the most common DNA binding motifs being AP-1, Grh and Sd (Figure 4A and Table S1). Strikingly, 78% (1,614 genes) of shared Yki/Sd target genes were also bound by Jun, indicating that in growing eyes, Yki, Sd and Jun bind to a largely overlapping set of target genes (Figure 4A). Consistent with this, the most enriched DNA binding motifs within shared Yki/Sd/Jun genome binding regions were AP-1, TEAD-Sd and Grh (Figure 4A). Ontology analyses revealed that the most enriched classes of genes bound by Yki/Sd/Jun belonged to the Hippo, MAPK and Wnt pathways (Figure 4B). Shared target genes included the previously identified Yki/Sd target genes *Diap1*, *ex*, *ban* and *kibra* and the JNK pathway genes *puckered* (*puc*), *matrix metalloprotease 1* (*Mmp1*), *cheerio* (*cher*) and *unpaired3* (*upd3*) ^3^ (Figures 4C and 4D). Many genes were also uniquely bound by either Yki/Sd or Jun, or not bound by any of these proteins but still expressed in larval eye discs (Figures S2D-S2F). Interestingly, previous ChIP-sequencing studies of the human Yki and Sd orthologues, YAP and TEAD, also reported enrichment of AP-1 motifs and functional relationships between YAP/TEAD and AP-1 were subsequently reported in cultured cell experiments ^36–38^. Similarly, the AP-1 motif was enriched in genes that were elevated in *warts* (*wts*) mutant *D. melanogaster* wing imaginal discs, which have elevated Yki activity ^39^. Therefore, our TaDa data, in conjunction with published genomic studies, indicate that the JNK and Hippo pathways share an overlapping transcriptome in both *D. melanogaster* and mammals. These findings, together with our genetic studies led us to postulate that JNK induction limits neoplastic growth by stimulating AP-1 to repress transcription of shared Hippo/JNK pathway genes.

**Figure 4.**
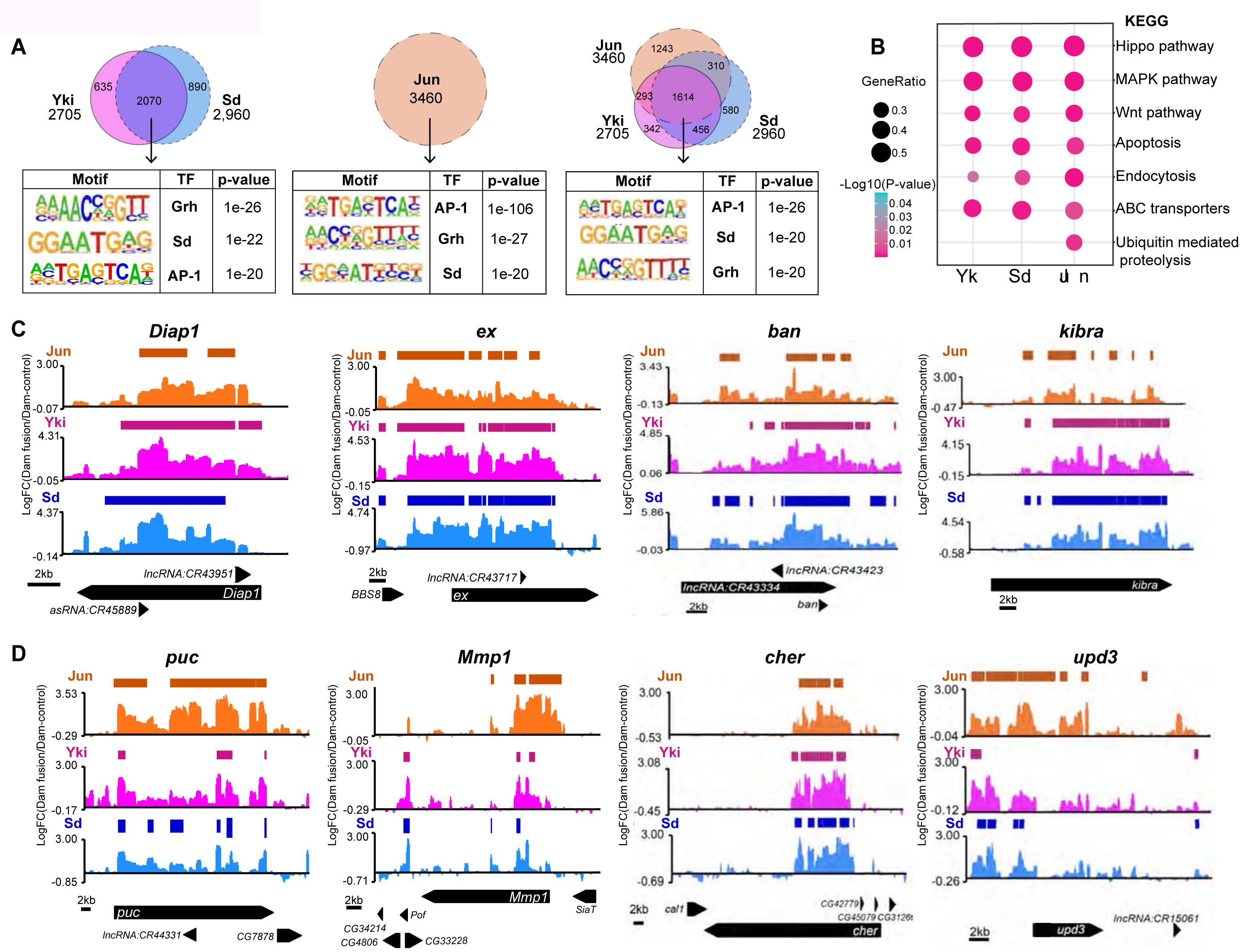
Hippo and JNK pathway transcription factors share a high degree of target genes in growing epithelial tissues. **A.** Venn diagrams showing the overlap of target genes bound by Yki, Sd and Jun. The AP-1, Sd/TEAD and Grh transcription factor motifs were the most enriched in shared Yki, Sd, and Jun target peaks. Motifs were identified using HOMER analysis with resampling 100 times using permutation (False Discovery Rate of 0.01). The significance associated with each motif is represented by a p-value. **B.** A bubble chart showing enrichment of KEGG pathways among target genes of Yki, Sd, and Jun as identified by TaDa. The KEGG pathways listed are statistically significant according to a p value of <0.05 to <0.001. **C-D.** Genome-binding profiles of target genes of Jun, Yki and Sd, as determined by DamID, for the Hippo pathway target genes *Diap1, ex, ban* and *kibra* (C), and the JNK/TNF pathway target genes *puc*, *Mmp1, cher* and *upd3* (D). Scale bars in kb are shown.

### The CtBP co-repressor and Fos repress transcription and neoplastic tumour growth

To investigate the mechanism by which AP-1 represses the transcription of growth genes in clonal neoplasia, we considered two transcriptional repressors: the Polycomb group and the C-terminal binding protein (CtBP). Loss of different Polycomb group genes induces dramatic overgrowth of *D. melanogaster* imaginal discs ^55,56^, and Polycomb group members influence neoplastic tissue growth and transcription in imaginal discs from homoygous *scrib* larvae ^44^. *CtBP* loss causes very mild overgrowth of imaginal discs, while CtBP and certain isoforms of Fos repress transcription of *ban*, which is a target of both Yki/Sd and AP-1 ^57^. To test potential roles for Polycomb group genes and CtBP in repression of transcription in *scrib* clones, we first used genetic mosaic experiments.

Depletion of the Polycomb group gene *Su(z)2* by RNAi does not obviously affect imaginal disc growth ^55^, but its depletion in hypomorphic *discs large* imaginal discs increases their growth ^44^. However, *Su(z)2* RNAi did not rescue the undergrowth of *scrib* clones in larval eye discs (Figures S3B, S3D and S3F). Additionally, RNAi-mediated depletion of the PRC1 component Polycomb (Pc) only modestly restored *scrib* clone growth, suggesting that Polycomb complex genes play a minor role in regulating the transcription of growth genes in clonal neoplasia (Figures S3B, S3C and S3F). Strikingly, however, RNAi-mediated depletion of CtBP completely restored the growth potential of *scrib* clones (Figures S3B, S3E and S3F). We tested this further by generating animals harbouring mutations in both *CtBP* and *scrib* on the FRT82B chromosome and assessed growth of larval eye-antennal imaginal discs by generating clones from the beginning of eye growth using *eyFlp*. *CtBP^87De10^, scrib*^1^ double mutant clones exhibited extremely strong overgrowth, similar to *scrib*^1^, *fos^KM^* eye clones, although more dramatic, with eye discs also becoming multi-layered and displaying clear apicobasal cell polarity defects (Figures 5A-5D, and Figure S4). In addition, *CtBP^87De10^, scrib*^1^ clones (in both eye-antennal and wing discs, generating using *hsFlp*) exhibited increased *ex-lacZ* levels, whilst single mutant *CtBP* clones and *scrib* clones did not display consistent changes in *ex-lacZ* (Figures 5E-5G’, S5G and S5K-L’). This latter result is consistent with a published study, which reported variable expression of Yki/Sd target genes in *scrib* clones ^9^. In addition, *DIAP1-lacZ*, *brC12*-lacZ and *ban3-GFP* were all elevated in *CtBP^87De10^, scrib*^1^ clones (Figures 5H-J’, S5C, S5F and S5H-J). *DIAP1-lacZ* and *ban3-GFP* were unchanged in single mutant clones, whilst *brC12*-lacZ was unchanged in *scrib*^1^ clones and mildly elevated in *CtBP^87De10^* clones (Figures 5E-5I’, S5A, B, D, E and S5H-J), consistent with a study that assessed *ban CtBP* wing imaginal disc clones ^57^.

**Figure 5.**
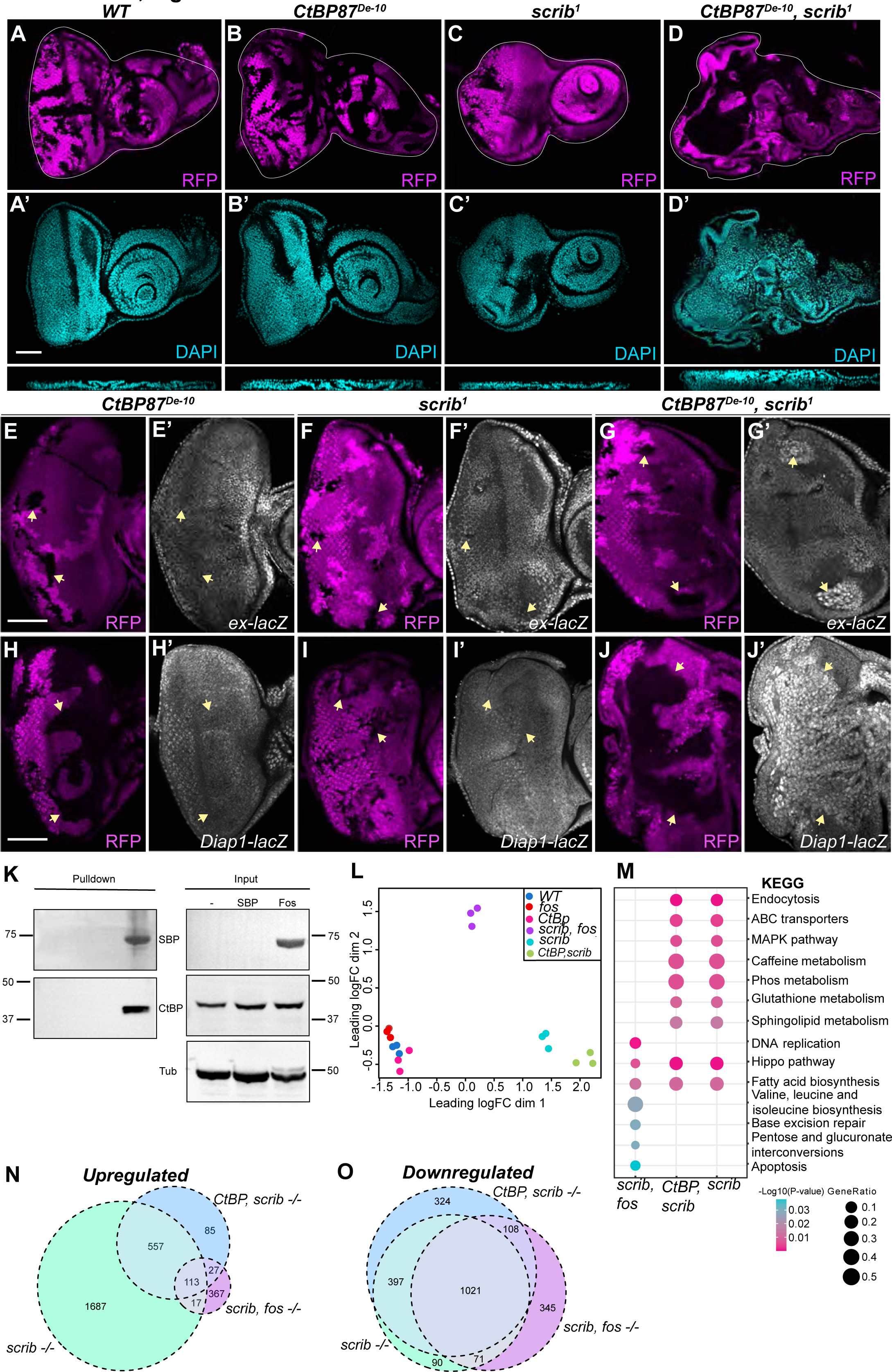
CtBP limits neoplastic clone growth by repressing Hippo and JNK pathway target genes. **A-D’.** Mosaic third instar larval eye-antennal discs. RFP (magenta) marks control wild-type tissue while non-RFP tissue is either wild-type (A) or mutant for the following genotypes: *CtBP^87De10^* (B), *scrib*^1^ (C), or *CtBP^87De10^, scrib*^1^ (D). DAPI marks nuclei (cyan in A’-D”) and below these XY images are XZ cross section images of these tissues. Clones were generated using *eyFlp*. The scale bar represents 50μm. **E-J’.** Mosaic third instar larval eye-antennal discs containing clones marked by the absence of RFP (magenta) of the following genotypes: *CtBP^87De10^* (E and H), *scrib*^1^ (F and I), or *CtBP^87De10^, scrib*^1^ (G and J). Tissues were stained with anti-β-Gal to reveal *ex-LacZ* expression (grayscale in E’-G’) and *Diap1-LacZ* expression (grayscale in I’-J’). Arrows indicate selected clones, which were generated using *hsFlp* in E-G’ and *eyFlp* in H-J’. Scale bars represent 50μm. **K.** Immunoblots of Fos-SBP pulldowns in S2 cells and input lysates, using the indicated antibodies. Molecular mass markers (kDa) are indicated. **L.** Multidimensional scaling plot of gene expression in third instar larval eye-antennal discs of the indicated genotypes. The x and y axes show the leading LogFC dimension 1 and 2 respectively. The two axes represent the variation in the data along two principal components. **M.** A bubble chart showing enrichment of KEGG pathways in genes that were differentially expressed in third instar larval eye-antennal discs of the following genotypes: *scrib^1^, fos^KM^; CtBP^87De10^, scrib*^1^; and *scrib*^1^. The KEGG pathways listed were statistically significant according to a p value of <0.05 to <0.001. **N-O.** Venn diagrams showing the degree of overlap of differentially expressed genes (upregulated in K and downregulated in L) in third instar larval eye-antennal discs of the following genotypes: *scrib*^1^*, fos^KM^; CtBP^87De10^, scrib*^1^; and *scrib*^1^.

These experiments provide strong evidence that CtBP represses transcription and growth of neoplastic clones together with AP-1 transcription factors. CtBP represses transcription by binding to transcription factors via a short consensus motif consisting of PxDLS, where x is any amino acid ^58,59^. Such a motif is present in Fos, but not Jun, Yki or Sd. Therefore, we assessed whether CtBP and Fos form a physical complex, by generating *D. melanogaster* S2 cell lines that stably expressed inducible plasmids encoding a Streptavidin Binding Protein (SBP)-tagged Fos protein or SBP alone. Endogenous CtBP protein was readily detected in SBP-Fos purifications by western blotting, but not in control SBP alone purifications, indicating that Fos and CtBP can indeed form a physical complex (Figure 5K). To investigate this further, we used RNA-seq to profile the transcriptome of late third instar larval eye imaginal discs harbouring predominantly *CtBP^87De10^* tissue or *CtBP^87De10^, scrib*^1^ double mutant tissue. *CtBP^87De10^* eyes had minor transcriptional changes compared to wild-type eyes, consistent with the fact that *CtBP* eye tissue has only a very mild growth advantage and normal tissue morphology ^57^ (Figures 5B and 5L). By contrast, the transcriptome of *CtBP^87De10^, scrib*^1^ eyes differed substantially from either control eyes or *CtBP^87De10^* eyes (Figures 5L). KEGG pathway and GO analysis revealed that genes belonging to the MAPK and Hippo pathways were highly elevated in *CtBP^87De10^, scrib*^1^ eyes, as were genes involved in endocytosis, metabolism, and actin binding (Figures 5M and S6A). This indicates that CtBP is a major repressor of transcription in neoplastic clones, in accordance with our genetic experiments and analysis of *ex, DIAP1 and ban* transcription reporters.

We performed similar experiments to investigate the role of Fos in transcription regulation in neoplastic clones. The transcriptome of eyes predominantly harbouring *scrib*^1^*, fos^KM^* tissue differed substantially from either control or *fos^KM^* eyes (Figure 5L), consistent with our genetic analyses. The most significantly elevated pathways and processes in *scrib*^1^*, fos^KM^* eyes were the Hippo pathway, DNA replication and fatty acid biosynthesis (Figure 5M). By comparing differentially expressed genes in *CtBP^87De10^, scrib*^1^ eyes and *scrib*^1^*, fos^KM^* eyes, we found significant overlaps of 140 upregulated genes (p<0.001, hypergeometric test) (Figure 5N), and 1,129 downregulated genes (p<0.001, hypergeometric test) (Figure 5O), consistent with the fact that loss of either *fos* or *CtBP* can repress the undergrowth of *scrib* clones. Finally, we compared the gene expression profile of *CtBP^87De10^, scrib*^1^ eyes and *scrib*^1^*, fos^KM^* eyes to *scrib*^1^ homozygous eye tissue. In a mosaic setting, *scrib* cells are eliminated, so we generated *scrib*^1^ eyes that were almost completely comprised of mutant tissue by flipping the *scrib*^1^ allele over a cell lethal mutation, which causes these tissues to overgrow strongly. The gene expression profile of *scrib*^1^ eye discs differed substantially from control eyes and, interestingly, overlapped considerably with the transcriptome of *CtBP^87De10^, scrib*^1^ eyes (2,088 shared differentially expressed genes) and *scrib*^1^*, fos^KM^* eyes (1,222 shared differentially expressed genes), further indicating that AP-1 and CtBP are major mediators of gene expression in *scrib* clones and hence their growth potential (Figures 5L-5O and S6A-S6C).

### Fos has both growth-repressive and growth-promoting potential in neoplastic cells

When examining gene expression changes in *CtBP^87De10^, scrib*^1^ and *scrib*^1^*, fos^KM^* eyes, we noticed that whilst genotypes both had a similar number of downregulated genes (1650 and 1332, respectively), the expression of many more genes were elevated in *CtBP^87De10^, scrib*^1^ tissue (1,002) than *scrib*^1^*, fos^KM^* tissue (203) (Figures 6A and 6B and Table S2). Multiple Hippo pathway genes were also elevated in homozygous *scrib* larvae eye discs, such as *ex, kibra* and *Mer* (Figure 6C), which contrasts with their being no change in *ex* abundance in *scrib* clones (Figure 2M). Presumably this is because in the mosaic scenario in (Figures 2M-P) JNK pathway induction supersedes the ability of Yki to induce target genes like *ex*, but in fully *scrib* mutant tissues (as in Figure 6C), JNK is not induced and Yki is free to activate its target genes.

**Figure 6.**
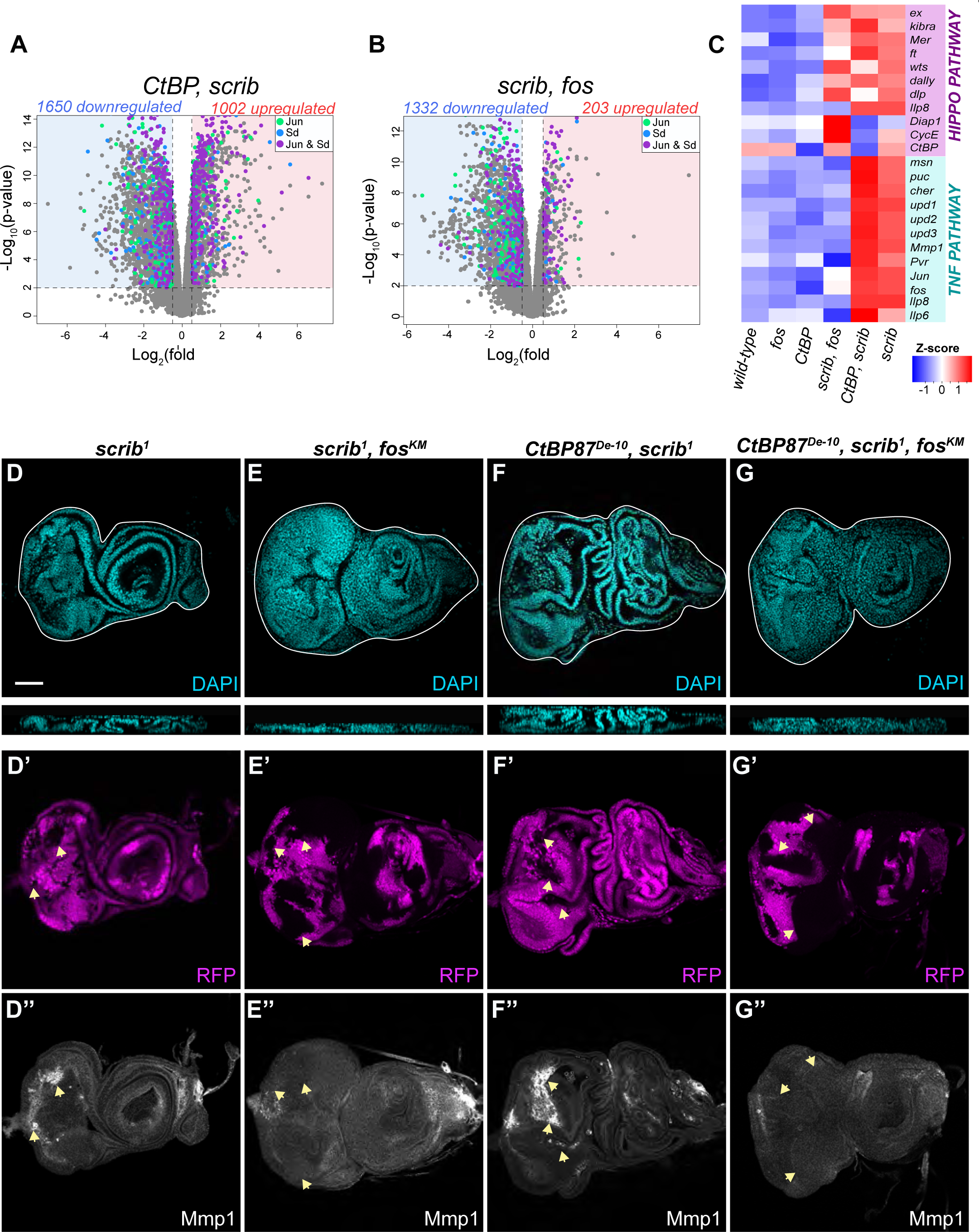
Fos has both growth-repressive and growth-promoting potential in neoplastic cells. **A-B.** Volcano plots displaying genes with upregulated expression (red box) or downregulated expression (blue) in *scrib*^1^*, fos^KM^* eye-antennal discs compared to *wild-type* discs (A), and *CtBP^87De10^, scrib*^1^ eye-antennal discs compared to wild-type discs (B). Samples shown in each coloured box had a fold change equal to or larger than 0.5 and an adjusted p-value of less than 0.01. Genes that were identified as candidate target genes using TaDa are highlighted in green for Jun, blue for Sd, and purple for Jun/Sd shared genes. **C.** A heatmap showing gene expression levels for known Hippo pathway target genes and known TNF pathway target genes. **D-G’’.** Mosaic third instar larval eye-antennal discs, DAPI marks nuclei (cyan in D-G). Below these XY images are XZ cross section images of these tissues. RFP (magenta in D’-G’) marks control wild-type tissue while non-RFP tissue is mutant for the following genotypes: *scrib* (D), *scrib*^1^*, fos^KM^* (E), *CtBP^87De10^, scrib*^1^ (F) or *CtBP^87De10^, scrib*^1^*, fos^KM^* (G). Mmp1 expression (grayscale in D’’-G’’) was revealed using an antibody. Arrows indicate selected clones, the scale bar represents 50μm.

Among the genes that were strongly elevated in *CtBP^87De10^, scrib*^1^ eyes but not *scrib, fos* eyes, were many JNK pathway genes such as *puc, cher, Mmp1, jun, fos* and *msn*, many of which are induced by AP-1 in feedback loops ^3^ (Figure 6C). We confirmed this by assessing expression of Mmp1, which is an AP-1 target that is induced in *scrib* clones by Fos ^10^. Mmp1 was strongly induced in both *scrib* clones and *CtBP^87De10^, scrib*^1^ clones but was not induced in *scrib*^1^*, fos^KM^* clones, consistent with our RNA-seq analyses (Figures 6D-6F”). This suggests that, upon JNK induction in *scrib* clones, Fos both activates and represses many genes, and this is why *CtBP^87De10^, scrib*^1^ eyes overgrow to a greater extent than *scrib*^1^*, fos^KM^* eyes (i.e., only Fos-dependent transcription repression is defective in *CtBP^87De10^, scrib*^1^ eyes whilst both Fos-mediated transcription repression and activation are lost in *scrib*^1^*, fos^KM^* eyes). Alternatively, CtBP loss might lead to greater overgrowth of *scrib* clones because it represses transcription and tissue growth through multiple transcription factors in addition to Fos. Indeed, CtBP has been reported to mediate transcription repression in the *Drosophila* eye via several other transcription factors ^60,61^.

To discern between these two possibilities, we generated *CtBP^87De10^, scrib*^1^*, fos^KM^* triple mutant animals and assessed the growth characteristics of clones of eye tissue mutant for all three genes. Eyes harbouring triple mutant eye clones closely resembled the phenotype of *scrib*^1^*, fos^KM^* eyes, as opposed to *CtBP^87De10^, scrib*^1^ eyes, as although *CtBP^87De10^, scrib*^1^*, fos^KM^* clones overgrew substantially, the eye discs did not become multi-layered (Figures 6D-6G). In addition, *CtBP^87De10^, scrib*^1^*, fos^KM^* triple mutant clones did express Mmp1 (Figure 6G-6G’’). Therefore, the fact that *CtBP^87De10^, scrib*^1^ eyes overgrow more than *scrib*^1^*, fos^KM^* eyes argues in favour of the first scenario, i.e., in clonal neoplasia, JNK induces Fos and CtBP to repress genes that are essential for clone growth and survival, but also induces Fos to activate a distinct set of genes, including those in the JNK pathway. This latter set of genes can also influence tissue growth, as well as apicobasal cell polarity, but this is only revealed when CtBP function is perturbed, and neoplastic clones are allowed to grow (Figure 7K). Notably, AP-1’s role in eye growth regulation was only apparent upon JNK pathway activation in the context of clonal neoplasia, as loss of *jun* or *fos* had no impact on eye growth in different growth scenarios when epithelial polarity was intact (normal growth, Yki-mediated hyperplasia or undergrowth of *yki* mutant tissue) (Figures S1D-S1F and Figure S7A-T).

**Figure 7.**
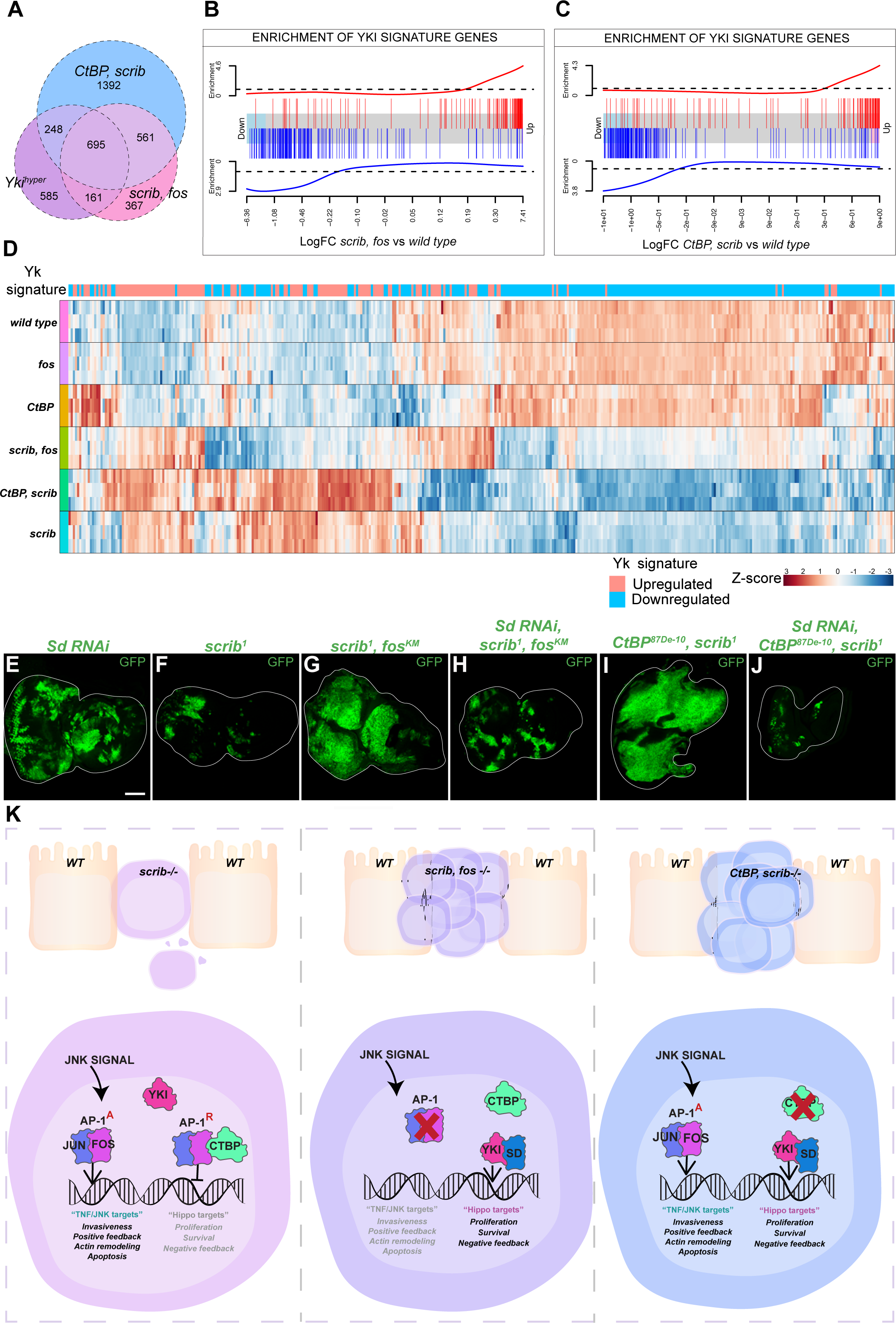
Yorkie and Scalloped drives the overgrowth of neoplastic tissues when AP-1/CtBP-mediated transcription repression is disabled. **A.** A Venn diagram showing the degree of overlap of differentially expressed genes in third instar larval eye-antennal discs of the following genotypes: *warts (*Yki hyperactive); *scrib*^1^*, fos^KM^;* and *CtBP^87De10^, scrib*^1^. **B-C.** Gene set enrichment plots showing enrichment of the Yki signature in differentially expressed genes in third instar larval eye-antennal discs of the following genotypes: *scrib*^1^*, fos^KM^* (B) and *CtBP^87De10^, scrib*^1^ (C). **D.** Heatmap showing expression levels of Yki signature genes in in third instar larval eye-antennal discs of the indicated genotypes (three replicates per genotype). **E-J.** Mosaic third instar larval eye-antennal discs harbouring eyFLP-MARCM-induced clones of the following genotypes: *Sd RNAi* (E), *scrib*^1^ (F), *scrib*^1^*, fos^KM^* (G), *Sd RNAi; scrib*^1^*, fos^KM^* (H), *CtBP^87De10^, scrib*^1^ (I), and *Sd RNAi; CtBP^87De10^, scrib*^1^ (H). MARCM clones are marked by GFP (green). The scale bars represents 50μm. **K.** A Schematic diagram showing AP-1 activating (AP-1^A^) and repressing roles (AP-1^R^) in response to induction of TNF signalling in neoplastic *scrib* clones. TNF/JNK signalling in activated in *scrib* clones, resulting in two different outcomes: 1) AP-1^A^ activates classic TNF target genes which promote apoptosis, actin remodeling and invasiveness, and 2) AP-1^R^, together with CtBP, represses Hippo pathway target genes, which can influence cell proliferation and viability.

### Yorkie drives the overgrowth of neoplastic tissues when Fos/CtBP-mediated transcription repression is disabled

Finally, we investigated the mechanism by which alleviation of AP-1/CtBP transcription repression in *scrib* clones facilitates the dramatic change of clone growth potential, i.e., from growth retardation to substantial overgrowth. Yki was a prime candidate for this given that we discovered it to be more nuclear in *scrib* clones and active in *scrib*^1^*, fos^KM^* and *CtBP^87De10^, scrib*^1^ clones, and because previous studies found that Yki activity was elevated when Scrib was depleted in broad epithelial tissue domains by RNAi ^62^. To investigate this, we performed RNA-seq of *wts* larval eye discs and found that differentially expressed genes strongly overlapped with gene expression changes in both *CtBP^87De10^, scrib*^1^ and *scrib*^1^*, fos^KM^* eyes (Figure 7A). We then compared genes that were either upregulated or downregulated in *wts* tissue with those that were bound by Yki and Sd in our TaDa experiments to identify a transcriptional signature of Yki hyperactivity (Table S3). The Yki hyperactive signature strongly correlated with both upregulated and downregulated genes in both *CtBP^87De10^, scrib*^1^ and *scrib*^1^*, fos^KM^* eye tissue (Figures 7B and 7C). This correlation was also evident when we performed unsupervised hierarchical clustering on the expression of Yki hyperactive signature genes in *CtBP, scrib* and *scrib*^1^*, fos^KM^* eyes, as well as wild-type, and *fos^KM^* and *Ctbp^87De10^* single mutant eyes (Figure 7D). Whilst wild-type eyes and *Ctbp^87De10^* and *fos^KM^* single mutant eyes all had very similar expression levels of Yki hyperactive signature genes, *CtBP^87De10^, scrib*^1^ and *scrib*^1^*, fos^KM^* displayed strong and highly overlapping changes in expression of these genes (Figure 7D, Table S4 and searchable web interface ^63^). Consistent with these gene expression studies, RNAi-mediated depletion of Sd strongly suppressed the overgrowth of both *scrib*^1^*, fos^KM^* clones and *CtBP^87De10^, scrib*^1^ clones (Figures 7E-7J).

## DISCUSSION

Polarized epithelia are essential features of most animal organs. Epithelial integrity is specified and maintained by a range of proteins including those that regulate apicobasal polarity and cell-cell adhesion ^1^. In addition, signalling pathways such as TNF/JNK and Hippo are important for sensing damaged epithelial cells and stimulating their removal from the epithelium ^3^. To date, most studies on this topic have focused on upstream signalling events that initiate removal of epithelial cells with defective apicobasal polarity caused by mutations in genes such as *scrib*. Many studies have reported that removal of defective epithelial cells is mediated by cell competition, a phenomenon where cells with a relative fitness advantage (“winner” cells) can actively outcompete “loser” cells^6^. However, a recent study challenged this view by showing that epithelial cells are removed by cell-autonomous TNF/JNK signalling ^7^. In this model, the TNF ligand is sequestered from its apically-localized receptor in cells with normal polarity, and TNF/JNK pathway activity is only induced in cells with defective polarity where the TNF receptor moves basally and can meets its ligand ^7^. In the present study, we provide important mechanistic insights into the role that control of transcription plays in the maintenance of epithelial integrity and prevention of tumour initiation.

Important roles for the JNK and Hippo pathways in epithelial growth control have been reported in numerous studies, although the relationships between these pathways have been somewhat contentious. For example, JNK has been reported to limit the growth of neoplastic epithelial clones by activating the Hippo pathway kinase Wts to repress its key effector, the transcription co-activator Yki ^35^. In contrast, the JNK pathway has also been reported to activate Yki in the context of wing regeneration, in this case by phosphorylating Jub, and increasing its ability to inhibit Wts ^34^. Here, we present several pieces of evidence that argue that JNK combats clonal neoplasia by acting via its downstream transcription factors, rather than by biochemical crosstalk with one or more Hippo pathway proteins. First, the Jun and Fos transcription factors are essential for the removal of defective epithelial cells from the eye (Figure 1) and ^10^. Second, TaDa studies in the growing eye revealed that transcription factors of the Hippo and JNK pathways share a largely overlapping suite of target genes (approximately 75% of Yki/Sd target genes are also bound by Jun) (Figure 4). Third, the Hippo pathway is not activated upon JNK pathway stimulation in *scrib* cells; rather Hippo activity is compromised in neoplastic clones, as Yki is substantially more nuclear and active (Figures 2, 3 and 7). The latter finding is supported by zebrafish and mammalian studies where Scrib promoted Hippo pathway activity ^64,65^, and by *D. melanogaster* studies where Yki was required for the overgrowth of homozygous *scrib* tissues ^62^.

Interestingly, both *jun* and *fos* were dispensable for normal eye growth as well as hyperplasia caused by Hippo pathway perturbation, and their growth regulatory potential was only revealed in the context of epithelial polarity loss. In this setting, our results argue that upon JNK induction in *scrib* cells, Fos represses transcription of shared JNK/Hippo pathway target genes to limit the growth potential of these cells and ensure their safe removal from the epithelium (Figure 7K). In support of this, we found that loss of the CtBP corepressor protein also dramatically reversed the growth potential of *scrib* cells, and loss of either *fos* or *CtBP* in *scrib* cells caused changes in expression of many of the same genes. Thus, we hypothesize that to limit the growth potential of epithelial cells that have lost polarity, JNK stimulates Fos and CtBP to modulate the chromatin environment of growth genes, rendering them impervious to activation by Yki and Sd (Figure 7K). Our genetic and RNA-seq analyses indicate that CtBP is a major regulator of gene expression in *scrib* clones, although additional transcription repressor complexes could also possibly be required for this. One such repressor complex that we considered is the Polycomb group complex, although we found only a minor role for it in limiting the growth potential of *scrib* clones. In future studies, it will be important to define the mechanism by which CtBP regulates the chromatin status of growth genes in neoplastic cells, as it is known to mediate three different enzymatic activities: histone deacetylation, di-methylation of histone H3K9 and demethylation of histone H3K4 ^59^.

Our finding that AP-1, Yki and Sd share many target genes in growing *Drosophila* eyes is consistent with genomics experiments in both *Drosophila* epithelial tissues and human cultured cells ^36–39,41,66^. As such, the regulation of an overlapping set of target genes by the JNK and Hippo pathways, many of which are linked to cell proliferation and survival, is also likely to be highly relevant for human tumorigenesis. Indeed, AP-1 and Yki/Sd-YAP/TEAD cooperatively regulate transcription in *D. melanogaster* tumours and human cancer cell lines and promote several cancer related phenotypes, such as cell survival and proliferation, invasion and metastasis, and drug resistance ^36–38,67,68^. It remains to be determined whether AP-1 can also antagonize YAP/TEAD’s oncogenic potential and ability to activate transcription in human cells, which could represent an important defence against tumour initiation.

## MATERIALS AND METHODS

### D. melanogaster husbandry

*D. melanogaster* strains and experimental crosses were maintained on *D. melanogaster* medium: 4.2% (w/v) yeast, 3.51% (w/v) glucose, 0.34% (w/v) agar, 3.98% (w/v) polenta 1.17% (w/v) Nipagin/Bavistan (10% p-hydroxybenzoic acid methyl ester C8H8O3). *D. melanogaster* strains were stored at room temperature (22°C) or at 18°C and experimental crosses were carried out at 25°C unless indicated otherwise. Imaginal disc clones were generated using either the FLP/FRT recombination system (*hsFlp* or *eyFlp) or* the mosaic analysis with a repressible marker (MARCM) system. Details of all strains are provided in the Supplementary Materials.

### CRISPR/Cas9 generation of mutant D. melanogaster

CRISPR/Cas9 was used to generate *fos* and *jun* mutant alleles, according to^69,70^. gRNA oligonucleotides were designed to target all *fos* or all *jun* isoforms. gRNA oligonucleotides were annealed, phosphorylated, and cloned into the pBFv-U6.2 plasmid. Transgenes were injected and integrated into the *attP40* or *attP2* sites using phiC31 integrase by BestGene (Chino Hills, CA). Transgenic gRNA strains were crossed to *nanos-cas9,* and individual founder males crossed to female balancer flies. Offspring were first tested for failure to complement deficiency strains covering either the *fos* or *jun* gene loci, then genetic lesions identified by Sanger sequencing. The *fos^KM^* allele has a 28 base pair deletion (deletion: 3R:29791441-29791469), resulting in a frameshift after amino acid 304 then a premature stop codon after 53 amino acids of a different coding sequence caused by the frameshift. The *jun^KM^* allele has a 2 base pair deletion (deletion: 2R:10097408-10097409), resulting in a frameshift after amino acid 54 then a premature stop codon after 7 amino acids of a different coding sequence caused by the frameshift.

### Immunostaining and microscopy

Imaginal discs from wandering third instar larvae were dissected into PBS on ice and fixed for 20 minutes in 4% (v/v) paraformaldehyde (PFA) at room temperature. Tissues were washed three times with 0.3% Triton X-100 in PBS (PBS/T), incubated with primary antibodies in 0.3% PBT with 10% Normal Goat Serum (NGS) overnight at 4°C or for 4 hours at room temperature, then washed three times in 0.3% PBT. Tissues were incubated with secondary antibodies conjugated to Alexa488, Alexa555, Alexa568 or Alex533 (Molecular Probes, ThermoFisher Scientific) at 1:500 in 0.3% PBT with 10% NGS and incubated either overnight at 4°C or for 2-4 hours at room temperature. Tissues were washed three times for 15 minutes in 0.3% PBS/T, incubated in mounting medium (90% glycerol in PBS) and mounted for imaging. Staged embryos were dechorionated for 1 minute with thin bleach, washed with distilled water and fixed on a shaker for 20 minutes in a 1:1 mix of 4% Paraformaldehyde (Proscitech) in 1xPBS and Heptane (Sigma). Embryos were then washed 3 times with absolute methanol and 3 times with PBS/T and blocked for 1 hour in 0.1% PBS/T containing 10% normal goat serum (NGS). They were then incubated overnight at 4°C with primary antibody (rat-anti DCAD2, DSHB, AB_528120) diluted 1:50 in 0.1% PBS/T containing 10% NGS, washed 3 times with 0.3% PBS/T, blocked for 1 hour in 0.1% PBS/T containing 10% NGS, and incubated for 2 hours at room temperature with secondary antibody (goat-anti rat, AlexaFluor 568, Thermo, AB_2534121) diluted 1:500 in 0.1% PBST containing 10% NGS. Embryos were washed 3 times in 0.1% PBS/T containing 10% NGS and stored in Vectashield prior to mounting in a glass depression slide dorsal side up. Details of antibodies and stains are provided in Supplementary Materials. Imaginal disc images were captured with an inverted Olympus FV3000 confocal microscope and embryo with an inverted Nikon C1 confocal microscope. Data analysis was performed using Fiji (ImageJ) and Adobe Photoshop (23.1.1 Release).

### Quantification and statistical analyses

To measure clone volume, fluorescence intensity was measured from three-dimensional 2.5µm spaced confocal Z stacks using Volocity software (Improvision). Reporter fluorescence quantification and Yki localisation analysis was performed using ImageJ/Fiji. For reporter fluorescence quantification, an ROI was selected spanning wild type and mutant clones, and mean intensities were measured on a representative single z-slice. For nuclear vs cytoplasmic localisation of Yki, segmentation of nuclei area and cytoplasmic area was performed on a single z-slice, using DAPI fluorescence to specify nuclei. Statistical analyses were carried out using GraphPad Prism 9. Details of statistical tests performed are listed in figure legends. All *n* numbers represent biological replicates.

## Targeted DamID

### A) Cloning of DamID fusion plasmids

To create a fusion protein with Dam positioned at the C terminus of Yki, a *Yki-Dam* fusion gene was synthesised by Biomatik. This sequence contained the full coding sequence of *yki*, followed by a myc tag, a short linker and then the *Dam* sequence. This cDNA was sub-cloned into the pUAST-attB-LT3-NDam plasmid using Kpn1 so that the Dam sequence was replaced by *yki-Dam*. The *Jun-Dam* fusion gene was created by amplifying the *jun* coding sequence, which was inserted into the multiple cloning site upstream of the *Dam* sequence.

### B) *D. melanogaster* genetics and staging

*Eyeless-Gal4 Gal80ts* was crossed to *UAS-LT3-Dam*, *UAS-LT3-Dam-Sd*, *UAS-LT3-Yki-Dam*, and *UAS-LT3-Dam-Jun*. Crosses were reared at 25°C and left to lay eggs over a 4-hour period. The eggs were then transferred to 18°C (restrictive temperature) for 4 days to repress the activity of the *Gal4*. Next, second instar larvae were transferred to 29°C (permissive temperature) for 24 hours, which induced the expression of the Dam fusion proteins for a defined developmental period. The eye-antennal imaginal discs and mouth-hooks of early third instar larvae were dissected into PBS on ice and then used for DNA extraction. Early third instar larvae were selected based on developmental timing and morphological features that distinguish them from other instar larvae stages. Dissected tissues were transferred to 100μL of TENS buffer (100mM Tris (pH 8.0), 5mM EDTA, 200mM NaCl, 0.2% SDS) on ice. The tissue samples were either stored at −80°C until further processed or used immediately. Approximately 300-400 eye-antennal imaginal discs were dissected for each genotype per experiment, and this was carried out in triplicate (n=3).

### C) DNA isolation

DNA isolation of dissected tissues was adapted from^71^. Samples were treated with 2μL Proteinase K (20mg/mL) overnight at 56°C. The following day, RNA was digested at 37°C for 30 minutes using 0.5μL of RNase A (100 mg/ml). Samples were transferred to a 1.5mL spin phase lock gel light tube (5prime, cat. #2302800) and 100μL of phenol:chloroform:isoamylalchol was added, gently mixed and centrifuged for 5 minutes at 14000 rpm at room temperature. The upper phase was transferred to a fresh 1.5mL eppendorf tube and the DNA was precipitated using 1μL glycogen, 10μL 3M NaAc, and 300μL EtOH for 30 minutes at −80°C, followed by centrifuging for 30 minutes at 14,000 rpm at 4°C. DNA pellets were washed using 70% EtOH, centrifuged and briefly air-dried. The DNA pellet was dissolved using 10-25μL of 10mM Tris ph8 and incubated at room temperature for 2-4 hours. DNA concentration was measured using Thermo-Fisher NanoDrop 2000 Spectrophotometer and either stored at −80°C or processed immediately.

### D) DamID-sequencing

The targeted DamID protocol was adapted from published studies^50,71,72^. Briefly, 2μg of DNA was digested overnight at 37°C in 0.5μL DpnI (10 units) and 1μL of 10X CutSmart buffer (NEB). A control sample was incubated without DpnI (-DpnI). Following overnight digestion, 0.5μL of DpnI was added for an additional 1 hour incubation, followed by heat inactivation at 80°C for 20 minutes. An adaptor ligation mix was added to each digested DNA sample, which contained 6.2μL H2O, 0.8μL 50uM double stranded adaptor AdR (40pmol), 2μL 10X ligation buffer and 1μL of T4 DNA ligase (5U/μL). A control was included that contained no ligase (-Lig). The ligation was carried out for 2 hours at 16°C, and the reaction was inactivated by a 10-minute incubation at 65°C.

The samples were then digested with DpnII for 4 hours at room temperature, where 1μL DpnII buffer, 0.2μL DpnII (50U), and 28.8μL MQ was added to each sample. The resulting digested and ligated DNA was then amplified by PCR. The mPCR included adding 20μL 5X MyTaq Reaction Buffer, 2.5μL Primer AdR (50μM), 2μL MyTaq DNA Polymerase, and 50.5μL MqH2O to 25μL of DpnII digested DNA. The DNA was amplified using the following PCR program:

68°C 10 min (1 cycle)

94°C 30 s; 65°C 5 min; 68°C 15min (1 cycle)

94°C 30 s; 65°C 1 min; 68°C 10min (3 cycles)

94°C 30 s; 65°C 1 min; 68°C 2min (14 cycles)

68°C 5 min (1 cycle)

5μL of mePCR product was loaded on a 1% agarose gel. If a smear was observed in experimental samples but was not present in the negative control samples (-DpnI and -Lig), the samples were PCR-purified using a QIAGEN PCR purification kit. Each sample was eluted twice in 30μL of elution buffer. The concentration of the purified DNA was measured using a Qubit fluorometer (dsDNA High Sensitivity Assay Kit, Thermo Fisher). DamID samples were adjusted to contain a total of 350ng of DNA in 55μL of elution buffer. The DNA was sonicated using Covaris S2 instrument to generate an average DNA fragment size of ∼300bp (duty cycle = 10%, intensity = 5 cycles, bust = 200, time = 45 sec, power = ∼23W, bath temperature = ∼4°C). Sonicated samples were digested with 1uL of Alw1 at 37°C overnight to remove DamID adaptors. DNA was purified using Agencourt AMPure XP. End repair and A-tailing and adaptor ligation was carried out using the KAPA Hyper Prep Kit. The libraries were amplified using the KAPA HiFi HotStart ReadyMix according to manufacturer’s instructions (Kapa Biosystems). Libraries were quantified using a Tapestation Bioanalyzer 2100 DNA 1000 chip and sequenced on an Illumina NextSeq instrument, using 75 base paired end (PE75) reads and 20 million reads per sample. For each sample, 3 replicates were carried out and DNA-seq samples were sequenced in several batches using the same protocol.

### E) DamID-sequencing analysis

DamID-seq datasets were processed to remove remaining DamID ligation adaptors and sequencing adaptors using Cutadapt. Alignment to the *D. melanogaster* reference genome (Release 6; dm6) was carried out using the Subread package. FeatureCounts was used to count sequencing reads and assign them to GATC features (subsequently referred to as ‘tags’). The edgeR package in R was used to determine differentially methylated tags of the Dam-fusion compared to the Dam-alone control. edgeR was selected due to its ability to integrate biological replicates and correct for batch effects in the analysis. The data was filtered so that tags that have 0.5 counts per million (cpm) in at least 3 samples were included in the analysis. Normalization was carried out using the TMM method and multiple hypothesis testing correction was performed. Significantly differentially methylated tags were then assigned to peaks (using Python Version 3.7.6), which indicate regions of specific binding of the Dam-fusion proteins to DNA. Peaks were associated with the nearest gene within a 5kb distance to the closest transcription start site (TSS) using bedtools closest gene function. For an example of this analysis pipeline, as well as the peak calling script, see https://github.com/jibsch/DamID_Seq_Analysis.

### RNA-sequencing and analysis

*eyeless-FLP; FRT82B Cell lethal* was crossed to: *FRT82B* (control), *FRT82B fos^KM^*, *FRT82B scrib*^1^, *FRT82B CtBP^87De-10^*, *FRT82B fos^KM^ scrib*^1^, and *FRT82B CtBP^87De-10^ scrib*^1^. Crosses were kept at 25°C and wandering third instar larvae were dissected into PBS on ice and transferred to Trizol. RNA was extracted using Trizol according to manufacturer’s instructions. Libraries were quantified using a Tapestation Bioanalyzer 2100 DNA 1000 chip. RNA-sequencing was carried out on Illumina NextSeq instrument, using 75 base paired end (PE75) reads and 20 million reads per sample. RNA-seq samples were sequenced in the same batch and 3 replicates were carried out for each genotype. Alignment of the RNA-seq data to the *D. melanogaster* reference genome (Release 6; dm6) was performed using the Subread package and FeatureCounts was used to generate a counts table. RNA-seq data analysis was carried out according to the module designed by COMINE-Australia (GitHub: https://github.com/COMBINE-Australia). The count data was imported and analysed in RStudio using the edgeR and limma packages. Lowly expressed genes were filtered and genes that have 0.5 counts per million (cpm) in at least 3 samples were included in the analysis. TMM normalization was implemented to eliminate composition biases between libraries. Differential expression analysis was carried out using the voom function in limma. The empirical Bayes method was used to perform empirical Bayes shrinkage on the variances, estimate moderated t-statistics and generate p-values and log2-fold-change. We implemented a fold change threshold of larger than 0.5 and an adjusted p-value threshold less than 0.01. Packages and tools used in this manuscript are listed in Supplementary Materials.

### Bioinformatics Analysis

Packages and tools used in this manuscript are listed in Supplementary Materials. A hypergeometric test was implemented using the phyper function in R and a gene universe of 13601 to estimate the probably of genes overlapping by chance alone.

### KEGG pathway and GO enrichment analysis

KEGG pathway enrichment and Gene Ontology (GO) analyses were performed in RStudio. Entrez IDs were generated using the AnnotationDbi and org.Dm.eg.db packages. KEGG pathway analysis was perfromed on Entrez IDs using the kegga function from the limma package and GO analysis on Entrez IDs using the goana function from the limma package. KEGG and GO data were filtered to include significant terms (p<0.05); if terms were duplicated, only the most significant term was used. KEGG and GO were plotted in RStudio using the ggplot2 package and visualized on a log10(p-value) scale.

### Transcription factor motif enrichment analysis

Transcription factor motif enrichment analysis was performed using Hypergeometric Optimization of Motif EnRichment (HOMER) in Terminal. The findMotifsGenome.pl tool was used to identify *de novo* motifs enriched in the DamID-seq peak files compared to the background *D. melanogaster* genome (dm6). A false discovery rate (FDR) was calculated from 100 randomisations, and a motif was considered significant according to a p-value < 0.05, and a FDR (q-value) of < 0.05.

### Yki hyperactive signature

To identify a hyperactive Yki signature, we compared genes that were differentially expressed (either upregulated or downregulated) in *wts* tissue and that were also bound by Yki and Sd in our TaDa experiments.

### Heatmaps

Heatmaps for the RNA-seq dataset were created using the heatmap.2 function from the gplots package in RStudio. The genes of interest and associated log counts were subset and the matrix created from this was used to create the heatmaps with Z-scores. Z-scores were calculated by taking the expression of a gene and subtracting the mean across all samples and dividing by the standard deviation. The RColorBrewer package was used for the colour scheme.

## Data and software availability

Raw DamID-seq and RNA-seq has been deposited at GEO. Software is available at: https://github.com/jibsch/DamID_Seq_Analysis

### *D. melanogaster* S2 cell line generation

The *fos* cDNA (isoform A) was synthesised by Biomatik and cloned into pMK33-SBP-N ^73^, using SpeI and XhoI. To generate S2 cell lines stably expressing SBP-Fos or SBP alone, three million S2 cells were seeded in Schneider’s medium with 10% FBS and 1% penicillin/streptomycin and transfected with 1mg pMK33-based plasmid using Effectene (Qiagen). 300 mg/ml hygromycin was added 48 hours later and cells cultured for 1 month to select for plasmid-expressing cells.

### Protein affinity purification and immunoblotting

To purify SBP-tagged proteins, S2 cells were incubated overnight with 75 mM CuSO_4_ to induce protein expression, washed twice in ice-cold PBS, then lysed for 20 minutes in 50 mM Tris, pH 7.5, 5% glycerol, 0.2% IGEPAL, 1.5 mM MgCl_2_, 125 mM NaCl, 25 mM NaF, 1 mM Na_3_VO_4_, 1 mM DTT, and Complete protease inhibitors (Roche). Lysates were clarified by centrifugation and incubated with streptavidin beads (Pierce) for 4 hours at 4°C. Beads were washed with lysis buffer four times, subjected to SDS-PAGE and transferred to Immobilon membrane (Millipore).

Membranes were blocked in 5% milk in Tris-buffered saline with 0.1% Tween (TBS/T), incubated with primary antibodies specific for SBP or CtBP overnight at 4°C in blocking buffer, washed 3 times, incubated with HRP-coupled secondary antibodies for 1 hour at room temperature and washed 3 times. Membranes were incubated with ECL Prime (GE Healthcare) and detected using a Bio-Rad Chemidoc system.

## ACKNOWLEDGEMENTS

We thank members of the Harvey lab for discussions. We thank I. Hariharan, B. Hay, H. Richardson, the Bloomington *Drosophila* Stock Center, the Vienna *Drosophila* RNAi Center, the Australian *Drosophila* Research Support Facility (www.ozdros.com), and the Developmental Studies Hybridoma Bank for *D. melanogaster* stocks and antibodies. This research was supported by the Australian Research Council (DP180102044, DP190101743 and DP230101406). K.F.H was supported by a Senior Research Fellowship (1078220) and Investigator grant (1194467) from the National Health and Medical Research Council of Australia (NHMRC), K.A.M. and J.M.P. were supported by Australian Postgraduate Awards and A.T.P. was supported by an NHMRC Senior Research Fellowship (1116955) and the Lorenzo and Pamela Galli Medical Research Trust. The research benefitted by support from the Victorian State Government Operational Infrastructure Support and Australian Government NHMRC Independent Research Institute Infrastructure Support. We acknowledge the Peter Mac Centre for Advanced Histology and Microscopy and the Molecular Genomics Core Facility and support to them from the Peter MacCallum Cancer Foundation and the Australian Cancer Research Foundation.

## AUTHOR CONTRIBUTIONS

Conceptualization: KAM, JV, KFH

Investigation: KAM, JV, JS, JP, KFH

Funding acquisition: KFH, ATP

Supervision: KFH, JV, ATP

Writing – original draft: KFH, KAM

Writing – review & editing: KAM, JV, JS, JP, ATP, KFH

## COMPETING INTERESTS

The authors declare no competing interests.

## DATA AND MATERIALS AVAILABILITY

All data are available in the main text or the supplementary materials.

## Notes

### Competing Interest Statement

The authors have declared no competing interest.

## REFERENCES

1. St Johnston, D., and Sanson, B. (2011). Epithelial polarity and morphogenesis. Curr Opin Cell Biol 23, 540–546. 10.1016/j.ceb.2011.07.005.

2. Hariharan, I.K., and Bilder, D. (2006). Regulation of imaginal disc growth by tumor-suppressor genes in Drosophila. Annu Rev Genet 40, 335–361.

3. La Marca, J.E., and Richardson, H.E. (2020). Two-Faced: Roles of JNK Signalling During Tumourigenesis in the Drosophila Model. Front Cell Dev Biol 8, 42. 10.3389/fcell.2020.00042.

4. Kockel, L., Zeitlinger, J., Staszewski, L.M., Mlodzik, M., and Bohmann, D. (1998). Jun in Drosophila Development: Redundant and nonredundant functions and regulation by two MAPK signal transduction pathways. Genes and Development.

5. Noselli, S., and Agnes, F. (1999). Roles of the JNK signaling pathway in Drosophila morphogenesis. Curr Opin Genet Dev 9, 466–472. 10.1016/S0959-437X(99)80071-9.

6. Yamamoto, M., Ohsawa, S., Kunimasa, K., and Igaki, T. (2017). The ligand Sas and its receptor PTP10D drive tumour-suppressive cell competition. Nature 542, 246–250. 10.1038/nature21033.

7. de Vreede, G., Gerlach, S.U., and Bilder, D. (2022). Epithelial monitoring through ligand-receptor segregation ensures malignant cell elimination. Science 376, 297–301. 10.1126/science.abl4213.

8. Brumby, A.M., and Richardson, H.E. (2003). scribble mutants cooperate with oncogenic Ras or Notch to cause neoplastic overgrowth in Drosophila. Embo J 22, 5769–5779.

9. Chen, C.L., Schroeder, M.C., Kango-Singh, M., Tao, C., and Halder, G. (2012). Tumor suppression by cell competition through regulation of the Hippo pathway. Proc Natl Acad Sci U S A 109, 484–489. 10.1073/pnas.1113882109.

10. Uhlirova, M., and Bohmann, D. (2006). JNK- and Fos-regulated Mmp1 expression cooperates with Ras to induce invasive tumors in Drosophila. EMBO J 25, 5294–5304. 10.1038/sj.emboj.7601401.

11. Igaki, T., Pastor-Pareja, J.C., Aonuma, H., Miura, M., and Xu, T. (2009). Intrinsic tumor suppression and epithelial maintenance by endocytic activation of Eiger/TNF signaling in Drosophila. Dev Cell 16, 458–465. 10.1016/j.devcel.2009.01.002.

12. Gaspar, P., and Tapon, N. (2014). Sensing the local environment: actin architecture and Hippo signalling. Curr Opin Cell Biol 31C, 74-83. 10.1016/j.ceb.2014.09.003.

13. Zheng, Y., and Pan, D. (2019). The Hippo Signaling Pathway in Development and Disease. Dev Cell 50, 264–282. 10.1016/j.devcel.2019.06.003.

14. Halder, G., Dupont, S., and Piccolo, S. (2012). Transduction of mechanical and cytoskeletal cues by YAP and TAZ. Nat Rev Mol Cell Biol 13, 591–600. 10.1038/nrm3416.

15. Harvey, K.F., Zhang, X., and Thomas, D.M. (2013). The Hippo pathway and human cancer. Nat Rev Cancer 13, 246–257. 10.1038/nrc3458.

16. Pantalacci, S., Tapon, N., and Leopold, P. (2003). The Salvador partner Hippo promotes apoptosis and cell-cycle exit in Drosophila. Nat Cell Biol 5, 921–927.

17. Tapon, N., Harvey, K.F., Bell, D.W., Wahrer, D.C.R., Schiripo, T.A., Haber, D.A., and Hariharan, I.K. (2002). salvador promotes both cell cycle exit and apoptosis in Drosophila and is mutated in human cancer cell lines. Cell 110, 467–478. 10.1016/S0092-8674(02)00824-3.

18. Wu, S., Huang, J., Dong, J., and Pan, D. (2003). hippo encodes a Ste-20 family protein kinase that restricts cell proliferation and promotes apoptosis in conjunction with salvador and warts. Cell 114, 445–456.

19. Udan, R.S., Kango-Singh, M., Nolo, R., Tao, C., and Halder, G. (2003). Hippo promotes proliferation arrest and apoptosis in the Salvador/Warts pathway. Nat Cell Biol 5, 914–920.

20. Harvey, K.F., Pfleger, C.M., and Hariharan, I.K. (2003). The Drosophila Mst ortholog, hippo, restricts growth and cell proliferation and promotes apoptosis. Cell 114, 457–467.

21. Kango-Singh, M., Nolo, R., Tao, C., Verstreken, P., Hiesinger, P.R., Bellen, H.J., and Halder, G. (2002). Shar-pei mediates cell proliferation arrest during imaginal disc growth in Drosophila. Development 129, 5719–5730.

22. Jia, J., Zhang, W., Wang, B., Trinko, R., and Jiang, J. (2003). The Drosophila Ste20 family kinase dMST functions as a tumor suppressor by restricting cell proliferation and promoting apoptosis. Genes Dev 17, 2514–2519.

23. Justice, R.W., Zilian, O., Woods, D.F., Noll, M., and Bryant, P.J. (1995). The Drosophila tumor suppressor gene warts encodes a homolog of human myotonic dystrophy kinase and is required for the control of cell shape and proliferation. Genes Dev 9, 534–546.

24. Xu, T., Wang, W., Zhang, S., Stewart, R.A., and Yu, W. (1995). Identifying tumor suppressors in genetic mosaics: the Drosophila lats gene encodes a putative protein kinase. Development 121, 1053–1063.

25. Lai, Z.C., Wei, X., Shimizu, T., Ramos, E., Rohrbaugh, M., Nikolaidis, N., Ho, L.L., and Li, Y. (2005). Control of cell proliferation and apoptosis by mob as tumor suppressor, mats. Cell 120, 675–685.

26. Dong, J., Feldmann, G., Huang, J., Wu, S., Zhang, N., Comerford, S.A., Gayyed, M.F., Anders, R.A., Maitra, A., and Pan, D. (2007). Elucidation of a universal size-control mechanism in Drosophila and mammals. Cell 130, 1120–1133.

27. Oh, H., and Irvine, K.D. (2008). In vivo regulation of Yorkie phosphorylation and localization. Development 135, 1081–1088. 10.1242/dev.015255.

28. Manning, S.A., Dent, L.G., Kondo, S., Zhao, Z.W., Plachta, N., and Harvey, K.F. (2018). Dynamic Fluctuations in Subcellular Localization of the Hippo Pathway Effector Yorkie In Vivo. Curr Biol 28, 1651–1660 e1654. 10.1016/j.cub.2018.04.018.

29. Zhang, L., Ren, F., Zhang, Q., Chen, Y., Wang, B., and Jiang, J. (2008). The TEAD/TEF family of transcription factor Scalloped mediates Hippo signaling in organ size control. Dev Cell 14, 377–387.

30. Wu, S., Liu, Y., Zheng, Y., Dong, J., and Pan, D. (2008). The TEAD/TEF family protein Scalloped mediates transcriptional output of the Hippo growth-regulatory pathway. Dev Cell 14, 388–398.

31. Goulev, Y., Fauny, J.D., Gonzalez-Marti, B., Flagiello, D., Silber, J., and Zider, A. (2008). SCALLOPED interacts with YORKIE, the nuclear effector of the hippo tumor-suppressor pathway in Drosophila. Curr Biol 18, 435–441.

32. Zhao, B., Wei, X., Li, W., Udan, R.S., Yang, Q., Kim, J., Xie, J., Ikenoue, T., Yu, J., Li, L., et al. (2007). Inactivation of YAP oncoprotein by the Hippo pathway is involved in cell contact inhibition and tissue growth control. Genes Dev 21, 2747–2761.

33. Sun, G., and Irvine, K.D. (2011). Regulation of Hippo signaling by Jun kinase signaling during compensatory cell proliferation and regeneration, and in neoplastic tumors. Dev Biol 350, 139–151. 10.1016/j.ydbio.2010.11.036.

34. Sun, G., and Irvine, K.D. (2013). Ajuba family proteins link JNK to Hippo signaling. Science signaling 6, ra81. 10.1126/scisignal.2004324.

35. Enomoto, M., Kizawa, D., Ohsawa, S., and Igaki, T. (2015). JNK signaling is converted from anti- to pro-tumor pathway by Ras-mediated switch of Warts activity. Dev Biol 403, 162–171. 10.1016/j.ydbio.2015.05.001.

36. Zanconato, F., Forcato, M., Battilana, G., Azzolin, L., Quaranta, E., Bodega, B., Rosato, A., Bicciato, S., Cordenonsi, M., and Piccolo, S. (2015). Genome-wide association between YAP/TAZ/TEAD and AP-1 at enhancers drives oncogenic growth. Nat Cell Biol 17, 1218–1227. 10.1038/ncb3216.

37. Stein, C., Bardet, A.F., Roma, G., Bergling, S., Clay, I., Ruchti, A., Agarinis, C., Schmelzle, T., Bouwmeester, T., Schubeler, D., and Bauer, A. (2015). YAP1 Exerts Its Transcriptional Control via TEAD-Mediated Activation of Enhancers. PLoS genetics 11, e1005465. 10.1371/journal.pgen.1005465.

38. Liu, X., Li, H., Rajurkar, M., Li, Q., Cotton, J.L., Ou, J., Zhu, L.J., Goel, H.L., Mercurio, A.M., Park, J.S., et al. (2016). Tead and AP1 Coordinate Transcription and Motility. Cell Rep 14, 1169–1180. 10.1016/j.celrep.2015.12.104.

39. Pascual, J., Jacobs, J., Sansores-Garcia, L., Natarajan, M., Zeitlinger, J., Aerts, S., Halder, G., and Hamaratoglu, F. (2017). Hippo Reprograms the Transcriptional Response to Ras Signaling. Dev Cell 42, 667–680 e664. 10.1016/j.devcel.2017.08.013.

40. Koo, J.H., Plouffe, S.W., Meng, Z., Lee, D.H., Yang, D., Lim, D.S., Wang, C.Y., and Guan, K.L. (2020). Induction of AP-1 by YAP/TAZ contributes to cell proliferation and organ growth. Genes Dev 34, 72–86. 10.1101/gad.331546.119.

41. Verfaillie, A., Imrichova, H., Atak, Z.K., Dewaele, M., Rambow, F., Hulselmans, G., Christiaens, V., Svetlichnyy, D., Luciani, F., Van den Mooter, L., et al. (2015). Decoding the regulatory landscape of melanoma reveals TEADS as regulators of the invasive cell state. Nat Commun 6, 6683. 10.1038/ncomms7683.

42. Goode, D.K., Obier, N., Vijayabaskar, M.S., Lie, A.L.M., Lilly, A.J., Hannah, R., Lichtinger, M., Batta, K., Florkowska, M., Patel, R., et al. (2016). Dynamic Gene Regulatory Networks Drive Hematopoietic Specification and Differentiation. Dev Cell 36, 572–587. 10.1016/j.devcel.2016.01.024.

43. Schaub, C., Rose, M., and Frasch, M. (2019). Yorkie and JNK revert syncytial muscles into myoblasts during Org-1-dependent lineage reprogramming. J Cell Biol 218, 3572–3582. 10.1083/jcb.201905048.

44. Bunker, B.D., Nellimoottil, T.T., Boileau, R.M., Classen, A.K., and Bilder, D. (2015). The transcriptional response to tumorigenic polarity loss in Drosophila. Elife 4. 10.7554/eLife.03189.

45. Riesgo-Escovar, J.R., Jenni, M., Fritz, A., and Hafen, E. (1996). The Drosophila Jun-N-terminal kinase is required for cell morphogenesis but not for DJun-dependent cell fate specification in the eye. Genes Dev 10, 2759–2768. 10.1101/gad.10.21.2759.

46. Willsey, H.R., Zheng, X., Carlos Pastor-Pareja, J., Willsey, A.J., Beachy, P.A., and Xu, T. (2016). Localized JNK signaling regulates organ size during development. Elife 5. 10.7554/eLife.11491.

47. Oh, H., and Irvine, K.D. (2011). Cooperative regulation of growth by Yorkie and Mad through bantam. Dev Cell 20, 109–122. 10.1016/j.devcel.2010.12.002.

48. Matakatsu, H., and Blair, S.S. (2012). Separating planar cell polarity and Hippo pathway activities of the protocadherins Fat and Dachsous. Development 139, 1498–1508. 10.1242/dev.070367.

49. Yu, J., and Pan, D. (2018). Validating upstream regulators of Yorkie activity in Hippo signaling through scalloped-based genetic epistasis. Development 145. 10.1242/dev.157545.

50. Southall, T.D., Gold, K.S., Egger, B., Davidson, C.M., Caygill, E.E., Marshall, O.J., and Brand, A.H. (2013). Cell-type-specific profiling of gene expression and chromatin binding without cell isolation: assaying RNA Pol II occupancy in neural stem cells. Dev Cell 26, 101–112. 10.1016/j.devcel.2013.05.020.

51. Wolff, T., and Ready, D.F. (1991). The beginning of pattern formation in the Drosophila compound eye: The morphogenetic furrow and the second mitotic wave. Development.

52. Ikmi, A., Gaertner, B., Seidel, C., Srivastava, M., Zeitlinger, J., and Gibson, M.C. (2014). Molecular evolution of the Yap/Yorkie proto-oncogene and elucidation of its core transcriptional program. Mol Biol Evol 31, 1375–1390. 10.1093/molbev/msu071.

53. Kowalczyk, W., Romanelli, L., Atkins, M., Hillen, H., Bravo Gonzalez-Blas, C., Jacobs, J., Xie, J., Soheily, S., Verboven, E., Moya, I.M., et al. (2022). Hippo signaling instructs ectopic but not normal organ growth. Science 378, eabg3679. 10.1126/science.abg3679.

54. Heinz, S., Benner, C., Spann, N., Bertolino, E., Lin, Y.C., Laslo, P., Cheng, J.X., Murre, C., Singh, H., and Glass, C.K. (2010). Simple Combinations of Lineage-Determining Transcription Factors Prime cis-Regulatory Elements Required for Macrophage and B Cell Identities. Molecular Cell. Elsevier Inc.

55. Classen, A.K., Bunker, B.D., Harvey, K.F., Vaccari, T., and Bilder, D. (2009). A tumor suppressor activity of Drosophila Polycomb genes mediated by JAK-STAT signaling. Nat Genet 41, 1150–1155. ng.445 [pii] 10.1038/ng.445.

56. Martinez, A.M., Schuettengruber, B., Sakr, S., Janic, A., Gonzalez, C., and Cavalli, G. (2009). Polyhomeotic has a tumor suppressor activity mediated by repression of Notch signaling. Nat Genet 41, 1076–1082. 10.1038/ng.414.

57. Sumabat, T.M., Worley, M.I., Pellock, B.J., Justin A. Bosch, J.A., Hariharan, I.K. (2019). The transcriptional co-repressor CtBP is a negative regulator of growth that antagonizes the Yorkie and JNK/AP-1 pathways. bioRxiv. 10.1101/772533.

58. Turner, J., and Crossley, M. (2001). The CtBP family: enigmatic and enzymatic transcriptional co-repressors. Bioessays 23, 683–690. 10.1002/bies.1097.

59. Chinnadurai, G. (2007). Transcriptional regulation by C-terminal binding proteins. Int J Biochem Cell Biol 39, 1593–1607. 10.1016/j.biocel.2007.01.025.

60. Barolo, S., Stone, T., Bang, A.G., and Posakony, J.W. (2002). Default repression and Notch signaling: Hairless acts as an adaptor to recruit the corepressors Groucho and dCtBP to Suppressor of Hairless. Genes Dev 16, 1964–1976. 10.1101/gad.987402.

61. Hoang, C.Q., Burnett, M.E., and Curtiss, J. (2010). Drosophila CtBP regulates proliferation and differentiation of eye precursors and complexes with Eyeless, Dachshund, Dan, and Danr during eye and antennal development. Dev Dyn 239, 2367–2385. 10.1002/dvdy.22380.

62. Doggett, K., Grusche, F.A., Richardson, H.E., and Brumby, A.M. (2011). Loss of the Drosophila cell polarity regulator Scribbled promotes epithelial tissue overgrowth and cooperation with oncogenic Ras-Raf through impaired Hippo pathway signaling. BMC Dev Biol 11, 57. 10.1186/1471-213X-11-57.

63. Mitchell et al., S.R.-s.d.f.F.D.h.d.e.m.e.d.c.h.c.f.a.b.e.a.

64. Skouloudaki, K., Puetz, M., Simons, M., Courbard, J.R., Boehlke, C., Hartleben, B., Engel, C., Moeller, M.J., Englert, C., Bollig, F., et al. (2009). Scribble participates in Hippo signaling and is required for normal zebrafish pronephros development. Proc Natl Acad Sci U S A 106, 8579–8584. 0811691106 [pii] 10.1073/pnas.0811691106.

65. Cordenonsi, M., Zanconato, F., Azzolin, L., Forcato, M., Rosato, A., Frasson, C., Inui, M., Montagner, M., Parenti, A.R., Poletti, A., et al. (2011). The Hippo transducer TAZ confers cancer stem cell-related traits on breast cancer cells. Cell 147, 759–772. 10.1016/j.cell.2011.09.048.

66. Zhang, X., Yang, L., Szeto, P., Abali, G.K., Zhang, Y., Kulkarni, A., Amarasinghe, K., Li, J., Vergara, I.A., Molania, R., et al. (2020). The Hippo pathway oncoprotein YAP promotes melanoma cell invasion and spontaneous metastasis. Oncogene 39, 5267–5281. 10.1038/s41388-020-1362-9.

67. Battilana, G., Zanconato, F., and Piccolo, S. (2021). Mechanisms of YAP/TAZ transcriptional control. Cell Stress 5, 167–172. 10.15698/cst2021.11.258.

68. Atkins, M., Potier, D., Romanelli, L., Jacobs, J., Mach, J., Hamaratoglu, F., Aerts, S., and Halder, G. (2016). An Ectopic Network of Transcription Factors Regulated by Hippo Signaling Drives Growth and Invasion of a Malignant Tumor Model. Curr Biol 26, 2101–2113. 10.1016/j.cub.2016.06.035.

69. Kondo, S., and Ueda, R. (2013). Highly improved gene targeting by germline-specific Cas9 expression in Drosophila. Genetics 195, 715–721. 10.1534/genetics.113.156737.

70. Vissers, J.H.A., Dent, L.G., House, C.M., Kondo, S., and Harvey, K.F. (2020). Pits and CtBP Control Tissue Growth in Drosophila melanogaster with the Hippo Pathway Transcription Repressor Tgi. Genetics 215, 117–128. 10.1534/genetics.120.303147.

71. Vissers, J.H.A., Froldi, F., Schroder, J., Papenfuss, A.T., Cheng, L.Y., and Harvey, K.F. (2018). The Scalloped and Nerfin-1 Transcription Factors Cooperate to Maintain Neuronal Cell Fate. Cell Rep 25, 1561–1576 e1567. 10.1016/j.celrep.2018.10.038.

72. Marshall, O.J., Southall, T.D., Cheetham, S.W., and Brand, A.H. (2016). Cell-type-specific profiling of protein-DNA interactions without cell isolation using targeted DamID with next-generation sequencing. Nat Protoc 11, 1586–1598. 10.1038/nprot.2016.084.

73. Kyriakakis, P., Tipping, M., Abed, L., and Veraksa, A. (2008). Tandem affinity purification in Drosophila: the advantages of the GS-TAP system. Fly (Austin) 2, 229–235. 6669 [pii].

