## Supplemental data for "The JNK and Hippo pathways control epithelial integrity and prevent tumour initiation by regulating an overlapping transcriptome"

**Supplemental information**

#### Supplemental Methods

##### *Drosophila melanogaster* genotypes

**Figure 1**
1A: *y, w, ey-FLP ; FRT42D Ubi-mRFP.nls / FRT42D; FRT82B Ubi-mGFP.nls / FRT82B*
1B: *y, w, ey-FLP ; FRT42D Ubi-mRFP.nls / FRT42D; FRT82B Ubi-mGFP.nls / FRT82B scrib*$^{1}$
1C: *y, w, ey-FLP ; FRT42D Ubi-mRFP.nls / FRT42D jun*$^{KM}$*; FRT82B Ubi-mGFP.nls / FRT82B scrib*$^{1}$
1E: *y, w, ey-FLP.N ; + / + ; FRT82B Ubi-mRFP.nls / FRT82B*
1F: *y, w, eyFLP.N ; + / + ; FRT82B Ubi-mRFP.nls / FRT82B fos*$^{KM}$
1G: *y, w, eyFLP.N ; + / + ; FRT82B Ubi-mRFP.nls / FRT82B scrib*$^{1}$
1H: *y, w, eyFLP.N ; + / + ; FRT82B Ubi-mRFP.nls / FRT82B scrib,*$^{1}$ *fos*$^{KM}$

**Figure 2**
2A: *y, w, eyFLP.N ; + / + ; Diap1-lacZ FRT82B Ubi-mRFP.nls / FRT82B fos*$^{KM}$
2B: *y, w, eyFLP.N ; + / + ; Diap1-lacZ FRT82B Ubi-mRFP.nls / FRT82B scrib*$^{1}$
2C: *y, w, eyFLP.N ; + / + ; Diap1-lacZ FRT82B Ubi-mRFP.nls / FRT82B scrib,*$^{1}$ *fos*$^{KM}$
2D: *y, w, hsFLP ; ex-lacZ / + ; FRT82B Ubi-mRFP.nls / FRT82B fos*$^{KM}$
2E: *y, w, hsFLP ; ex-lacZ / + ; FRT82B Ubi-mRFP.nls / FRT82B scrib*$^{1}$
2F: *y, w, hsFLP ; ex-lacZ / + ; FRT82B Ubi-mRFP.nls / FRT82B scrib,*$^{1}$ *fos*$^{KM}$
2G: *y, w, eyFLP.N / ban-GFP ; + / + ; FRT82B Ubi-mRFP.nls / FRT82B fos*$^{KM}$
2H: *y, w, eyFLP.N / ban-GFP; + / + ; FRT82B Ubi-mRFP.nls / FRT82B scrib*$^{1}$
2I: *y, w, eyFLP.N / ban-GFP; + / + ; FRT82B Ubi-mRFP.nls / FRT82B scrib,*$^{1}$ *fos*$^{KM}$
2J: *y, w, eyFLP.N ; + / + ; brC12-lacZ FRT82B Ubi-mRFP.nls / FRT82B fos*$^{KM}$
2K: *y, w, eyFLP.N ; + / + ; brC12-lacZ FRT82B Ubi-mRFP.nls / FRT82B scrib*$^{1}$
2L: *y, w, eyFLP.N ; + / + ; brC12-lacZ FRT82B Ubi-mRFP.nls / FRT82B scrib,*$^{1}$ *fos*$^{KM}$

**Figure 3**
3A: *y, w, ey-FLP.N ; + / + ; FRT82B Ubi-mRFP.nls / FRT82B*
3B: *y, w, eyFLP.N ; + / + ; FRT82B Ubi-mRFP.nls / FRT82B fos*$^{KM}$
3C: *y, w, eyFLP.N ; + / + ; FRT82B Ubi-mRFP.nls / FRT82B scrib*$^{1}$
3D: *y, w, eyFLP.N ; + / + ; FRT82B Ubi-mRFP.nls / FRT82B scrib,*$^{1}$ *fos*$^{KM}$

**Figure 5**
5A: *y, w, ey-FLP.N ; + / + ; FRT82B Ubi-mRFP.nls / FRT82B*
5B: *y, w, eyFLP.N ; + / + ; FRT82B Ubi-mRFP.nls / FRT82B CtBP*$^{87De\text{-}10}$
5C: *y, w, eyFLP.N ; + / + ; FRT82B Ubi-mRFP.nls / FRT82B scrib*$^{1}$
5D: *y, w, eyFLP.N ; + / + ; FRT82B Ubi-mRFP.nls / FRT82B CtBP*$^{87De\text{-}10}$ *, scrib*$^{1}$
5E: *y, w, hsFLP ; ex-lacZ / + ; FRT82B Ubi-mRFP.nls / FRT82B CtBP*$^{87De\text{-}10}$
5F: *y, w, hsFLP ; ex-lacZ / + ; FRT82B Ubi-mRFP.nls / FRT82B scrib*$^{1}$
5G: *y, w, hsFLP ; ex-lacZ / + ; FRT82B Ubi-mRFP.nls / FRT82B CtBP*$^{87De\text{-}10}$ *, scrib*$^{1}$
5H: *y, w, eyFLP.N ; + / + ; Diap1-lacZ FRT82B Ubi-mRFP.nls / FRT82B CtBP*$^{87De\text{-}10}$
5I: *y, w, eyFLP.N ; + / + ; Diap1-lacZ FRT82B Ubi-mRFP.nls / FRT82B scrib*$^{1}$

5J: *y, w, eyFLP.N ; + / + ; Diap1-lacZ FRT82B Ubi-mRFP.nls / FRT82B CtBP*$^{87De-10}$ *, scrib*$^{1}$

**Figure 6**
6D: *y, w, eyFLP.N ; + / + ; FRT82B Ubi-mRFP.nls / FRT82B scrib*$^{1}$
6E: *y, w, eyFLP.N ; + / + ; FRT82B Ubi-mRFP.nls / FRT82B scrib*$^{1}$*, fos*$^{KM}$
6F: *y, w, eyFLP.N ; + / + ; FRT82B Ubi-mRFP.nls / FRT82B CtBP*$^{87De-10}$ *, scrib*$^{1}$
6G: *y, w, eyFLP.N ; + / + ; FRT82B Ubi-mRFP.nls / FRT82B CtBP*$^{87De-10}$ *, scrib*$^{1}$*, fos*$^{KM}$

**Figure 7**
6E: *y, w, eyFLP ; act>y+>GAL3, UAS-GFP / UAS-SdRNAi ; FRT82B tubGAL80 / FRT82B*
6F: *y, w, eyFLP ; act>y+>GAL3, UAS-GFP / + ; FRT82B tubGAL80 / FRT82B scrib*$^{1}$
6G: *y, w, eyFLP ; act>y+>GAL3, UAS-GFP / + ; FRT82B tubGAL80 / FRT82B scrib*$^{1}$*, fos*$^{KM}$
6H: *y, w, eyFLP ; act>y+>GAL3, UAS-GFP / UAS-SdRNAi ; FRT82B tubGAL80 / FRT82B scrib*$^{1}$*, fos*$^{KM}$
6I: *y, w, eyFLP ; act>y+>GAL3, UAS-GFP / + ; FRT82B tubGAL80 / FRT82B CtBP*$^{87De-10}$ *, scrib*$^{1}$
6J: *y, w, eyFLP ; act>y+>GAL3, UAS-GFP / UAS-SdRNAi ; FRT82B tubGAL80 / FRT82B CtBP*$^{87De-10}$ *, scrib*$^{1}$

**Supplemental Figure 1**
S1C: *w1118, fos*$^{KM}$ */ fos*$^{KM}$*, fos*$^{2}$ */ fos*$^{2}$*,jun*$^{KM}$ */ jun*$^{KM}$*, jun*$^{A109}$ */ jun*$^{A109}$
S1D: *y, w, ey-FLP ; FRT42D Ubi-mRFP.nls / FRT42D; FRT82B Ubi-mGFP.nls / FRT82B*
S1E: *y, w, ey-FLP ; FRT42D Ubi-mRFP.nls / FRT42D jun*$^{KM}$*; FRT82B Ubi-mGFP.nls / FRT82B fos*$^{KM}$
S1G: *y, w, ey-FLP.N ; + / + ; FRT82B Ubi-mRFP.nls / FRT82B*
S1G': *y, w, eyFLP.N ; + / + ; FRT82B Ubi-mRFP.nls / FRT82B fos*$^{2}$
S1G'': *y, w, eyFLP.N ; + / + ; FRT82B Ubi-mRFP.nls / FRT82B scrib*$^{1}$
S1G''': *y, w, eyFLP.N ; + / + ; FRT82B Ubi-mRFP.nls / FRT82B scrib,*$^{1}$ *fos*$^{2}$
S1H: *y, w, ey-FLP.N ; + / + ; FRT82B Ubi-mRFP.nls / FRT82B*
S1H': *y, w, hsFLP ; ex-lacZ / + ; FRT82B Ubi-mRFP.nls / FRT82B fos*$^{KM}$
S1H'': *y, w, hsFLP ; ex-lacZ / + ; FRT82B Ubi-mRFP.nls / FRT82B scrib*$^{1}$
S1H''': *y, w, hsFLP ; ex-lacZ / + ; FRT82B Ubi-mRFP.nls / FRT82B scrib,*$^{1}$ *fos*$^{KM}$
S1I: *y, w, eyFLP.N ; + / + ; FRT82B Ubi-mRFP.nls / FRT82B scrib*$^{2}$

**Supplemental Figure 3**
S3A: *y, w, eyFLP ; act>y+>GAL3, UAS-GFP / + ; FRT82B tubGAL80 / FRT82B*
S3B: *y, w, eyFLP ; act>y+>GAL3, UAS-GFP / + ; FRT82B tubGAL80 / FRT82B scrib*$^{1}$
S3C: *y, w, eyFLP ; act>y+>GAL3, UAS-GFP / UAS-Pc RNAi ; FRT82B tubGAL80 / FRT82B scrib*$^{1}$
S3D: *y, w, eyFLP ; act>y+>GAL3, UAS-GFP / UAS-Su(z)2 RNAi ; FRT82B tubGAL80 / FRT82B scrib*$^{1}$
S3E: *y, w, eyFLP ; act>y+>GAL3, UAS-GFP / UAS-CtBP RNAi ; FRT82B tubGAL80 / FRT82B scrib*$^{1}$

**Supplemental Figure 4**
S4A: *y, w, eyFLP.N ; + / + ; brC12-lacZ FRT82B Ubi-mRFP.nls / FRT82B CtBP*$^{87De-10}$

S4B: *y, w, eyFLP.N ; + / + ; brC12-lacZ FRT82B Ubi-mRFP.nls / FRT82B scrib*$^{1}$
S4C: *y, w, eyFLP.N ; + / + ; brC12-lacZ FRT82B Ubi-mRFP.nls / FRT82B CtBP*$^{87De\text{-}10}$*, scrib*$^{1}$
S4D: *y, w, eyFLP.N / ban-GFP ; + / + ; FRT82B Ubi-mRFP.nls / FRT82B CtBP*$^{87De\text{-}10}$
S4E: *y, w, eyFLP.N / ban-GFP; + / + ; FRT82B Ubi-mRFP.nls / FRT82B scrib*$^{1}$
S4F: *y, w, eyFLP.N / ban-GFP; + / + ; FRT82B Ubi-mRFP.nls / FRT82B CtBP*$^{87De\text{-}10}$*, scrib*$^{1}$
S4K: *y, w, hs-FLP ; exLacZ / + ; FRT82B Ubi-GFP / FRT82B scrib*$^{1}$
S4L: *y, w, hs-FLP ; exLacZ / + ; FRT82B Ubi-GFP / FRT82B CtBP*$^{87De\text{-}10}$*, scrib*$^{1}$

**Supplemental Figure 5**
S5A: *y, w, eyFLP.N ; + / + ; FRT82B Ubi-mRFP.nls / FRT82B*
S5B: *y, w, eyFLP.N ; + / + ; FRT82B Ubi-mRFP.nls / FRT82B scrib*$^{1}$
S5C: *y, w, eyFLP.N ; + / + ; FRT82B Ubi-mRFP.nls / FRT82B scrib*$^{1}$*, fos*$^{KM}$
S5D: *y, w, eyFLP.N ; + / + ; FRT82B Ubi-mRFP.nls / FRT82B CtBP*$^{87De\text{-}10}$*, scrib*$^{1}$

**Supplemental Figure 7**
S7A,E: *y, w, ey-FLP.N ; FRT42D Ubi-mRFP.nls / FRT42D*
S7B, F: *y, w, ey-FLP.N ; FRT42D Ubi-mRFP.nls / FRT42D jun*$^{KM}$
S7C, G: *y, w, ey-FLP.N ; FRT42D Ubi-mRFP.nls / FRT42D Hpo*$^{5.1}$
S7D, H: *y, w, ey-FLP.N ; FRT42D Ubi-mRFP.nls / FRT42D jun*$^{KM}$*, Hpo*$^{5.1}$
S7I, M: *y, w, ey-FLP.N ; + / + ; FRT82B Ubi-mRFP.nls / FRT82B*
S7J, N: *y, w, ey-FLP.N ; + / + ; FRT82B Ubi-mRFP.nls / FRT82B fos*$^{KM}$
S7K, O: *y, w, ey-FLP.N ; + / + ; FRT82B Ubi-mRFP.nls / FRT82B sav*$^{3}$
S7L, P: *y, w, ey-FLP.N ; + / + ; FRT82B Ubi-mRFP.nls / FRT82B sav*$^{3}$*, fos*$^{KM}$
S7Q: *y, w, ey-FLP.N ; FRT42D Ubi-mRFP.nls / FRT42D*
S7R: *y, w, ey-FLP.N ; FRT42D Ubi-mRFP.nls / FRT42D jun*$^{KM}$
S7S: *y, w, ey-FLP.N ; FRT42D Ubi-mRFP.nls / FRT42D yki*$^{B5}$
S7T: *y, w, ey-FLP.N ; FRT42D Ubi-mRFP.nls / FRT42D yki*$^{B5}$*, jun*$^{KM}$

| REAGENT or RESOURCE | SOURCE | IDENTIFIER |
|---|---|---|
| **Fly strains** | | |
| y, w, ey-FLP.N | Bloomington Drosophila Stock Center (BDSC) | 5580 |
| FRT82B Ubi-mRFP.nls | BDSC | 30555 |
| FRT82B | BDSC | 2035 |
| FRT82B fos$^{KM}$ | Harvey lab | N/A |
| FRT42D jun$^{KM}$ | Harvey lab | N/A |
| FRT82B scrib$^{1}$ | (Bilder and Perrimon, 2000) Gift from Helena Richardson | N/A |

| | | |
|---|---|---|
| FRT82B scrib$^{2}$ | (Bilder and Perrimon, 2000) Gift from Helena Richardson | N/A |
| CtBP$^{87De-10}$ | BDSC | 1663 |
| ex$^{697}$ | BDSC | 44248 |
| FRT82B Ubi-GFP | BDSC | 5188 |
| hs-FLP.D5 | BDSC | 55815 |
| UAS-Sd RNAi | Vienna Drosophila Resource Centre (VDRC) | KK 101497 |
| w;; p[pUAS-LT3-Dam] attP2/TM6B | (Southall et al. 2013; Vissers et al. 2018) | N/A |
| w;; p[pUAS-LT3-Dam-Sd] attP2/TM6B | (Southall et al. 2013; Vissers et al. 2018) | N/A |
| w;; p[pUAS-LT3-Yki-Dam] attP2/TM6B | This study | N/A |
| w;; p[pUAS-LT3-Dam-Jun] attP2/TM6B | This study | N/A |
| ey-Gal4 | BDSC | 5534 |
| tub-Gal80ts | BDSC | 7017 |
| GMR-Gal4 | BDSC | 9146 |
| Nanos-cas9 | BDSC | 54591 |
| y, w, eyFLP ; act>y+>GAL3, UAS-GFP; FRT82B tubGAL80 / TM6B | This study | N/A |
| FRT82B | BDSC | 2035 |
| FRT42D | BDSC | 1802 |
| *Sd$^{47M}$* | (Srivastava et al., 2004) | N/A |
| *banLacZ* | BDSC | 10154 |
| jra$^{IA109}$ | BDSC | 3273 |
| *Hpo5.1* | (Genevet et al., 2009) | N/A |
| *Sav$^{3}$* | (Tapon et al., 2002) | N/A |

| | | |
|---|---|---|
| $Yki^{B5}$ | (Huang et al., 2005) | N/A |
| UAS-Su(z)2 RNAi | BDSC | 57466 |
| UAS-Pc RNAi | BDSC | 36070 |
| UAS-CtBP RNAi | VDRC | 107313 |
| **Antibodies** | | |
| Mouse anti-Diap1 (1:200) | Bruce Hay | N/A |
| Mouse anti-βgalactosidase (1:100) | Sigma-Aldrich | Cat#G4644 |
| Mouse anti-Crumbs | Developmental Studies Hybridoma Bank (DSHB) | Cq4 |
| Rabbit anti-aPKC (1:500) | Santa Cruz Biotech | Sc-216 |
| Mouse anti-Mmp1 | DSHB | 3A6B4 |
| Guinea pig anti-Yki | Iswar Hariharan | N/A |
| Donkey anti-mouse 647 | ThermoFisher Scientific | Cat#A31571 |
| Goat anti-guinea pig 647 | ThermoFisher Scientific | Cat#A21450 |
| Goat anti-rabbit 488 | ThermoFisher Scientific | Caat#A11008 |
| **Chemicals, Peptides, and Recombinant Proteins** | | |
| DAPI (1:500) | Sigma-Aldrich | Cat#D9542 |
| Proteinase K (20mg/mL) | Sigma | Cat#E6779 |
| RNase A (100mg/mL) | Qiagen | Cat#19101 |
| Phenol:chloroform:isoamyl alcohol (25:24:1) | Sigma | Cat#P2069 |
| DpnI (10U) T4 | NEB | Cat#R0176L |
| T4 DNA ligase (5 U/µL) | Sigma | Cat#10799009001 |
| DpnII (50U) | NEB | Cat#R0543L |
| MyTaq HS DNA polymerase | Bioline | Cat# BIO-21112 |
| AlwI | NEB | Cat#R0513L |
| Agencourt AMPure XP Beads | BeckmanCoulter | Cat#A63881 |
| Bbs1 | NEB | Cat# R0539 |
| T4 Polynucleotide Kinase | NEB | Cat# M0201S |
| Antarctic Phosphatase | NEB | Cat# M0289S |

| | | |
|---|---|---|
| Kpn1 | NEB | Cat# R0142S |
| DH5alpha E.coli | Thermo Fisher Scientific | Cat#18265017 |
| TRIzol | Thermo Fisher Scientific | Cat#15596018 |
| **Critical Commercial Assays** | | |
| MinElute PCR Purification Kit | Qiagen | Cat#28004 |
| KAPA HyperPrep Kit | KAPA Biosystems | Cat#07962371001 |
| QIAquick Gel purification kit | Qiagen | Cat# 28706 |
| Miniprep Kit | Qiagen | Cat# 27106 |
| **Oligonucleotides** | | |
| AdRt: CTAATACGACTCACTATAGGG CAGCGTGGTCGCGGCC GAGGA | (Vogel, Peric-Hupkes, and van Steensel 2007) | |
| AdRb: TCCTCGGCCG | (Vogel, Peric-Hupkes, and van Steensel 2007) | |
| AdR_PCR: GGTCGCGGCCGAGGATC | (Vogel, Peric-Hupkes, and van Steensel 2007) | |
| fos crispr gRNA fwd: CTTC GACCACCACGCGCAACATCG | This study | |
| fos crispr gRNA rev:AAAC CGATGTTGCGCGTGGTGGTC | This study | |
| jun crispr gRNA fwd: CTTC GGGGTGGATGTGTTCAGGTT | This study | |
| jun crispr gRNA rev: AAAC AACCTGAACACATCCACCCC | This study | |
| **Recombinant DNA** | | |
| pBFv-U6.2 | (Kondo and Ueda 2013) | |
| pBFv-U6.2-fos gRNA | This study | |
| pBFv-U6.2-jun gRNA | This study | |
| pUASt-LT3-Dam | (Southall et al. 2013) | |
| pUASt-LT3-Dam Sd | (Vissers et al. 2018) | |

| | |
|---|---|
| pUASt-LT3-Yki Dam | This study |
| pUASt-LT3-Dam Jun | This study |
| **Software, packages and toolkits** | |
| AnnotationDbi (V1.48.0) | (Pag.s et al., 2019) |
| Bedtools (v2.28.0) | (Quinlan and Hall 2010) |
| call_peaks.py | This study |
| Cutadapt (v1.10) | (Martin 2011) |
| dplyr (v0.8.3) | (Wickham et al., 2019) |
| edgeR (v3.26.8) | (Robinson et al., 2010) |
| EnhancedVolcano (v1.2.0) | (Blighe, 2019) |
| eulerr (v6.0.0) | (Larsson, 2019) |
| FeatureCounts (v1.6.4) | (Liao et al., 2014) |
| ggplot2 (v3.2.1) | (Wickham, 2016) |
| gplots (v 3.0.1.1) | (Warnes et al., 2019) |
| Glimma (v1.12.0) | (Su et al., 2017) |
| GO.db (v3.8.2) | (Carlson, 2019) |
| gridExtra (v2.3) | (Auguie, 2017) |
| HOMER (v4.11) | (Heinz et al., 2010) |
| limma (v3.40.6) | (Richie et al., 2015) |
| KEGG.db (v3.2.3) | (Carlson, 2016) |
| org.Dm.eg.db (v3.8.2) | (Carlson, 2019) |
| Rprojroot (v1.3.2) | (Müller, 2018) |
| stringr (v1.4.0) | (Wickham, 2019) |
| Subread (v2.0.0) | (Liao et al., 2013) |
| svglite (v1.2.2) | (Wickham et al., 2019) |
| tidyr (v1.0.0) | (Wickham and Henry, 2019) |
| tidyverse (v1.2.1) | (Wickham, 2017) |

**Supplemental Figure Legends**

**Figure S1. The TNF/JNK pathway transcription factors Jun and Fos are dispensable for developmental eye growth, related to Figure 1.**

**A-B.** Schematic diagram of the *D. melanogaster fos* and *jun* loci and CRISPR/Cas9 genome editing of the *fos* and *jun* loci. Exons from each *fos* and *jun* isoforms are shown as purple boxes and introns are shown as black lines. The location and sequence of the gRNA target and PAM are shown. The deleted sequence for each gene is shown with dashed lines. In the bottom panel, the *wild-type* and mutant proteins are shown. The target site of the gRNA is shown in red, and the basic region and leucine zipper (bZIP) domain is shown in blue. The *fos* and *jun* mutations results in a frameshift that induces a premature STOP codon and a truncated protein.

**C-D.** Dorsal-view maximum-projection images of stage 14/15 *D. melanogaster* embryos of the indicated genotypes, immunostained for E-cadherin (greyscale). Anterior is up in all images. Scale bar corresponds to 20 µm.

**D-E.** Mosaic third instar larval eye-antennal discs containing clones marked by the absence of either RFP expression or GFP expression, for *wild-type* controls (*FRT42D*, *FRT82B*) (D), or $jun^{KM}$ and $fos^{KM}$ (E). The boxed regions are magnified below. Overlapping mutant clones are outlined in a dashed yellow line. Images shown represent single slice confocal micrographs. Scale bars correspond to 50µm.

**F.** Chart showing quantification of clone size in the genotypes shown in D-E, shown as a ratio of $RFP^-$ volume, or $GFP^-$ volume, over total eye-antennal disc volume. The genotypes are: *FRT42D* (shown in D), $jun^{KM}$ (shown in E), *FRT82B* (shown in D'), $fos^{KM}$ (shown in E'), overlapping *FRT42D* and *FRT82B* clones (shown D''), and overlapping $jun^{KM}$ and $fos^{KM}$ clones (shown in E''). n = 8. Data are represented as mean ± SEM. p-values were obtained using unpaired t-test, ns = not significant.

**G-I.** Mosaic third instar larval eye-antennal (G and I) and wing (H) discs containing clones of the indicated genotypes marked by the absence of either RFP or GFP expression. Expression of Diap1 (G), *ex-LacZ* (H) and Yki (I), all in greyscale, were revealed by antibodies. DAPI is cyan in I, arrows highlight select clones. Scale bars correspond to 50µm.

**Figure S2. Hippo and TNF/JNK pathway transcription factors share a high degree of target genes in growing epithelial tissues, related to Figure 4.**

**A.** Multidimensional scaling plot representation of DamID-seq samples. DamID-seq samples were plotted in two dimensions so that distances on the plot approximate differences between samples based on log2 fold changes. The x and y axes show the leading LogFC dimension 1 and 2 respectively. The plot displays each DamID-seq sequencing batch (shown as triangles, boxes, or circles), which are samples that were sequenced in separate sequencing runs, and the DamID-seq genotype (shown as different colours).

**B.** Density plots displaying the number of reads per peak for each replicate in the DamID-seq experiments.

**C.** Venn diagrams showing the degree of overlap between DamID-seq datasets and published datasets; Yki-Dam compared to Yki-ChIP (Ikmi et al., 2014), Sd-Dam compared to Sd-ChIP (Ikmi et al., 2014), Sd-Dam compared to Sd ChIP-Nexus (Kowalczyk et al., 2022), and Sd-ChIP (Ikmi et al., 2014) compared to Sd ChIP-Nexus (Kowalczyk et al., 2022).

**D-F.** Genome-binding profiles of Yki, Sd, and Jun as determined by DamID, for genes that are uniquely bound by Sd and Yki, but not Jun (D), genes that are uniquely bound by Jun, but not Sd and Yki (E), and genes that are highly expressed in the third instar larval eye-antennal disc, but not bound by Yki, Sd, and Jun (F).

**Figure S3. The Polycomb group is not a major mediator of repression of transcription or growth of neoplastic clones, related to Figure 5.**

**A-E.** Mosaic third instar larval eye-antennal discs of ey-FLP-MARCM-induced clones of *wild-type (A), scrib$^{1}$* (b), Pc RNAi, *scrib$^{1}$* (C), *Su(z)2 RNAi, scrib$^{1}$* (D), and *CtBP RNAi*, *scrib$^{1}$* (E). DAPI is cyan, and MARCM clones are marked by the presence of GFP. Images shown represent single slice confocal micrographs. Scale bars correspond to 50µm.

**F.** Chart showing quantification of size of clones of the genotypes in A-E, shown as a ratio of $GFP^{+}$ volume over total eye-antennal disc volume. The genotypes are: *FRT82B wildtype, FRT82B scrib$^{1}$*; Pcr RNAi, *scrib$^{1}$*; *Su(z)2 RNAi, scrib$^{1}$*; and *CtBP RNAi*, *scrib$^{1}$*. n = 8, 26, 6, 10, and 9, respectively. Data are represented as mean ± SEM. p-values were obtained using unpaired t-tests. ** $p < 0.01$, **** $p < 0.0001$.

**Figure S4. Fos and CtBP impact epithelial tissue architecture to different degrees, related to Figure 5.**

**A-D**. Mosaic third instar larval eye-antennal discs containing clones marked by the absence of RFP expression of *wild-type* (*WT*) (A), $scrib^{1}$ (B), $scrib^{1}, fos^{KM}$ (C), or $CtBP^{87De10}, scrib^{1}$ (D). Eye-antennal discs were stained for apical polarity markers using anti-aPKC (green) and anti-Crumbs (red). DAPI is shown in cyan. xy images show a single slice confocal micrograph of the apical region of the eye-antennal disc, and xz panels show cross sections. $scrib^{1}$ (B), $scrib^{1}, fos^{KM}$ (C), and $CtBP^{87De10}, scrib^{1}$ clones have perturbed localisation of the apical polarity proteins aPKC and Crumbs. Note that $CtBP^{87De10}, scrib^{1}$ (D) displays the most severe tissue architecture and epithelial polarity defects. Arrowheads indicate representative clones. Scale bars correspond to 50μm.

**Figure S5. CtBP limit neoplastic clone growth and expression of Hippo pathway target genes, related to Figure 5.**

**A-F'.** Mosaic third instar larval wing imaginal discs containing clones marked by the absence of RFP (magenta) of the following genotypes: $CtBP^{87De10}$ (A and D), $scrib^{1}$ (B and E), or $CtBP^{87De10}, scrib^{1}$ (C and F). Tissues were stained with anti-β-Gal to reveal *brC12-LacZ* expression (grayscale in A'-C') and *ban-GFP* is in grayscale (D-F'). Arrowheads indicate selected clones, the scale bar represents 50μm.

**G-J.** Charts showing quantification of *ex-lacZ* (G), *Diap1-lacZ* (H), *ban-GFP* (I) or *brC12-lacZ* (J) in mutant versus wild-type larval eye disc tissue of the indicated genotypes. n = 4, 4, 7, 6 in (G), 7, 7, 8, 5 in (H), 6, 5, 11, 7 in (I), and 6, 8, 6, 6 in (J). Data are represented as mean ± SEM. p-values were obtained using unpaired t-tests. * $p < 0.05$, ** $p < 0.01$, *** $p < 0.001$.

**K-L'.** Mosaic third instar larval wing imaginal discs containing clones marked by the absence of RFP (magenta) of the following genotypes: $scrib^{1}$ (K), or $CtBP^{87De10}, scrib^{1}$ (L). Tissues were stained with anti-β-Gal to reveal *ex-LacZ* expression (grayscale in K' and L'). Arrowheads indicate selected clones, the scale bar represents 50μm.

**Figure S6. Fos and CtBP regulate a largely overlapping transcriptome, related to Figure 6.**

**A.** A bubble chart showing enrichment of Gene Ontology (GO) terms performed on genes that have significantly upregulated expression in *scrib*$^{1}$*, fos*$^{KM}$; *CtBP*$^{87De10}$*, scrib*$^{1}$; and *scrib*$^{1}$ eye-antennal discs.
**B.** A bubble chart showing enrichment of KEGG pathways performed on genes that have significantly downregulated expression in *scrib*$^{1}$*, fos*$^{KM}$; *CtBP*$^{87De10}$*, scrib*$^{1}$; and *scrib*$^{1}$ eye-antennal discs.
**C.** A bubble chart showing enrichment of Gene Ontology (GO) terms performed on genes that have significantly downregulated expression in *scrib*$^{1}$*, fos*$^{KM}$; *CtBP*$^{87De10}$*, scrib*$^{1}$; and *scrib*$^{1}$ eye-antennal discs.

**Figure S7. AP-1 is dispensable for both hyperplasia caused by Hippo pathway perturbation and Hippo pathway-mediated default repression, related to Figure 6.**
**A-D.** Mosaic adult *Drosophila* female eyes containing clones marked by the absence of pigment and of the following genotypes: *wild-type* (*WT*) (A), *jun*$^{KM}$ (B), *hpo*$^{5.1}$ (C), or *jun*$^{KM}$*, hpo*$^{5.1}$ (D).
**E-F.** Mosaic eye-antennal discs containing clones marked by the absence of RFP expression of *wild-type* (*WT*) (E), *jun*$^{KM}$ (F), *hpo*$^{5.1}$ (G), or *jun*$^{KM}$*, hpo*$^{5.1}$ (H). Eye-antennal discs were stained with anti-Diap1 (grey), shown in bottom panel. Arrowheads indicate representative clones. Scale bars correspond to 50µm.
**I-L.** Mosaic adult *Drosophila* female eyes containing clones marked by the absence of pigment and of the following genotypes: *wild-type* (*WT*) (I), *jun*$^{KM}$ (J), *hpo*$^{5.1}$ (K), or *jun*$^{KM}$*, hpo*$^{5.1}$ (L).
**M-P.** Mosaic eye-antennal discs containing clones marked by the absence of RFP expression of *wild-type* (*WT*) (M), *fos*$^{KM}$ (N), *sav*$^{3}$ (O), or *sav*$^{3}$*, fos*$^{KM}$ (P). Eye-antennal discs were stained with anti-Diap1 (grey), shown in bottom panel. Arrowheads indicate representative clones. Scale bars correspond to 50µm.
**Q-T.** Loss of *jun* does not rescue *yki* clones, or Diap1 protein levels, in the eye-antennal disc. Mosaic eye-antennal discs containing clones marked by the absence of RFP expression of *wild-type* (*WT*) (Q), *jun*$^{KM}$ (R), *yki*$^{B5}$ (S), or *jun*$^{KM}$*, yki*$^{B5}$ (T). Eye-antennal discs were stained with anti-Diap1 (grey), shown in bottom panel. Images shown represent single slice confocal micrographs. Scale bars correspond to 50µm.

**Table S1. Yki, Sd and Jun target genes, related to Figure 4.**

Genes identified by targeted DamID as bound by Yki-Dam, Sd-Dam, and Jun-Dam.

**Table S2. Identification of genes that are responsive to Fos, CtBP and Scrib, related to Figures 6 and 7.**

Differentially expressed genes in larval eye-antennal discs. The following RNA-seq datasets were compared: *CtBP, scrib* and control; *scrib*, *fos* and control; and *scrib* and control.

**Table S3. Yki hyperactive target gene signature, related to Figure 7.**

Genes that were bound by Yki and Sd in targeted DamID studies and were either upregulated or downregulated in *warts* mutant eye-antennal discs, as determined by RNA-seq.

**Table S4. Differentially expressed genes in eye-antennal discs of different genotypes and Yki/Sd target gene status, related to Figure 7.**

Values represent log2 normalised RNA seq counts.

**Mitchell *et al.*, Fig. S1**

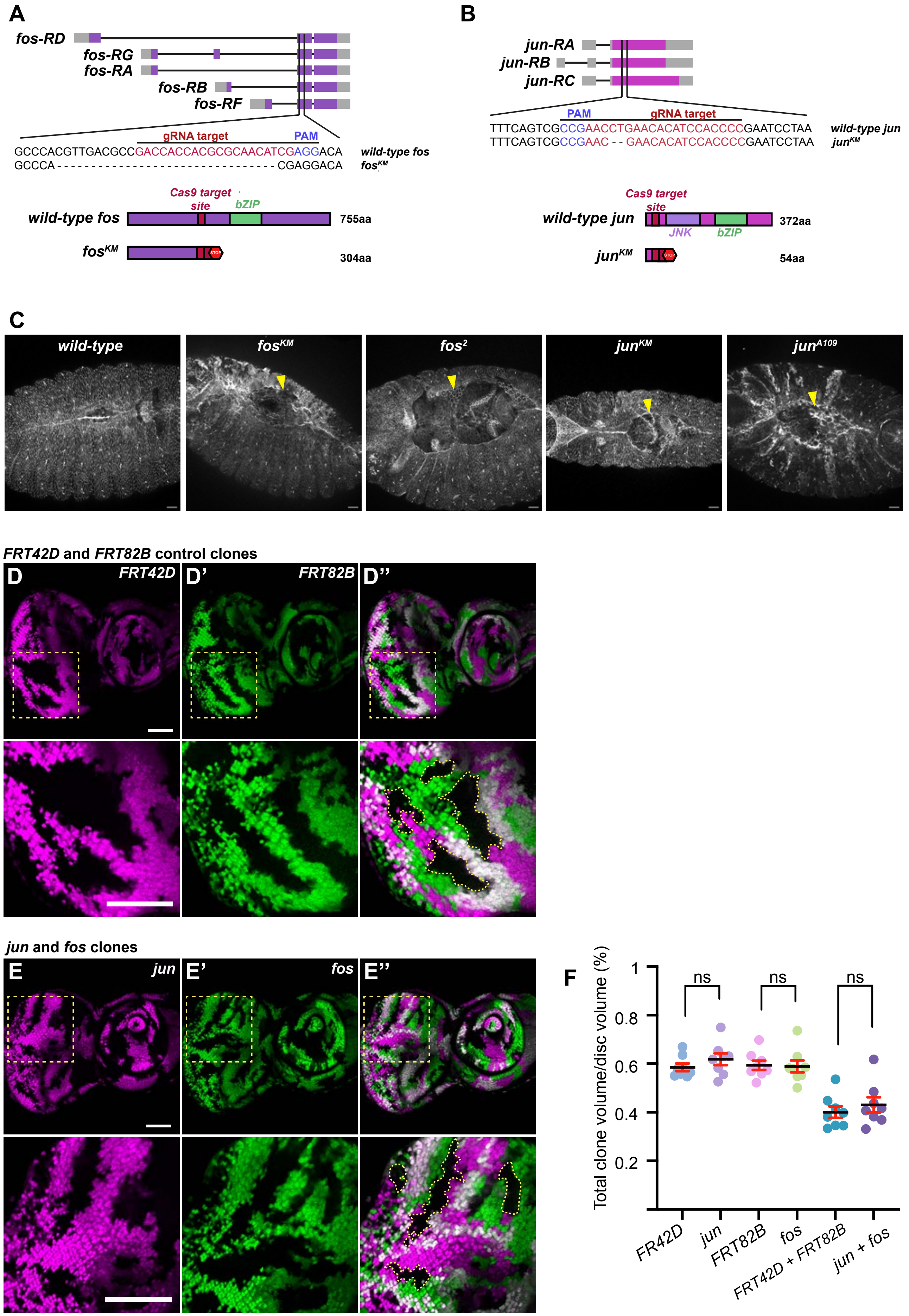

Mitchell *et al.*, Fig. S2

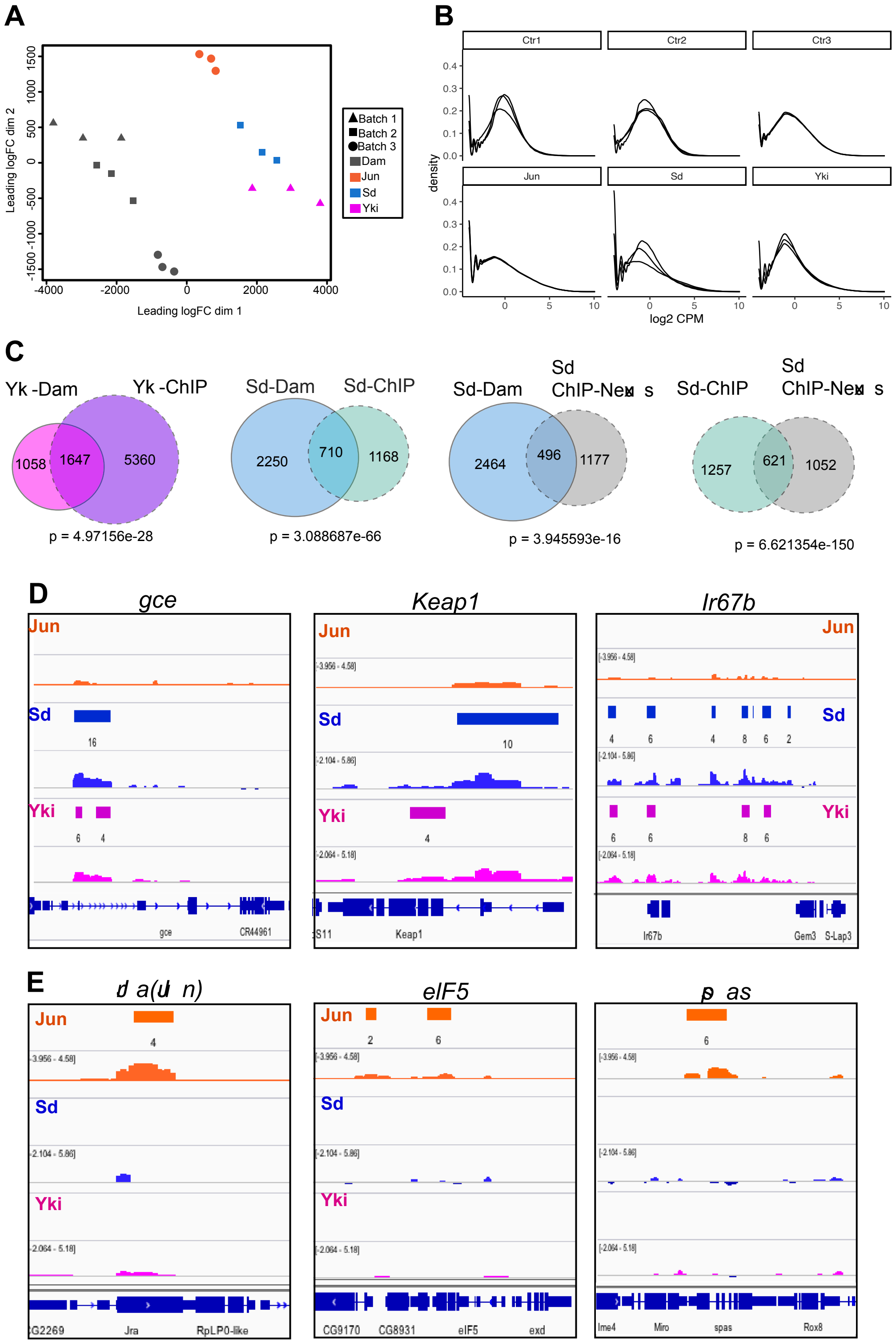

Mitchell *et al.*, Fig. S2

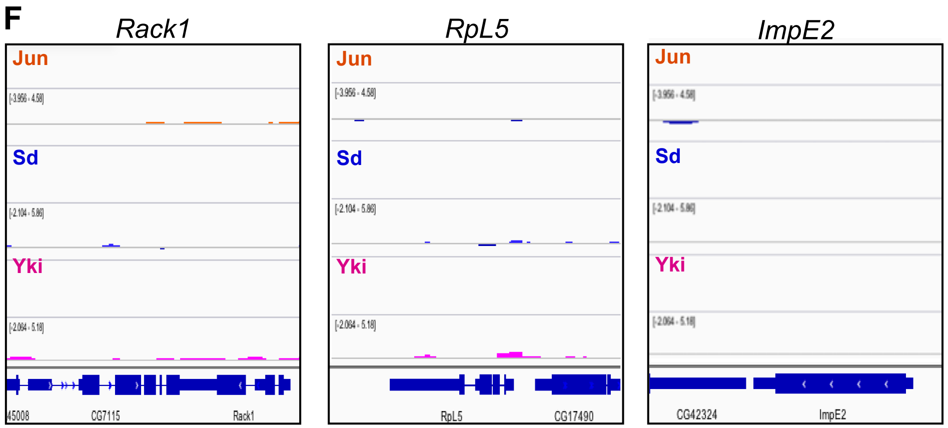

**Mitchell *et al*., Fig. S3**

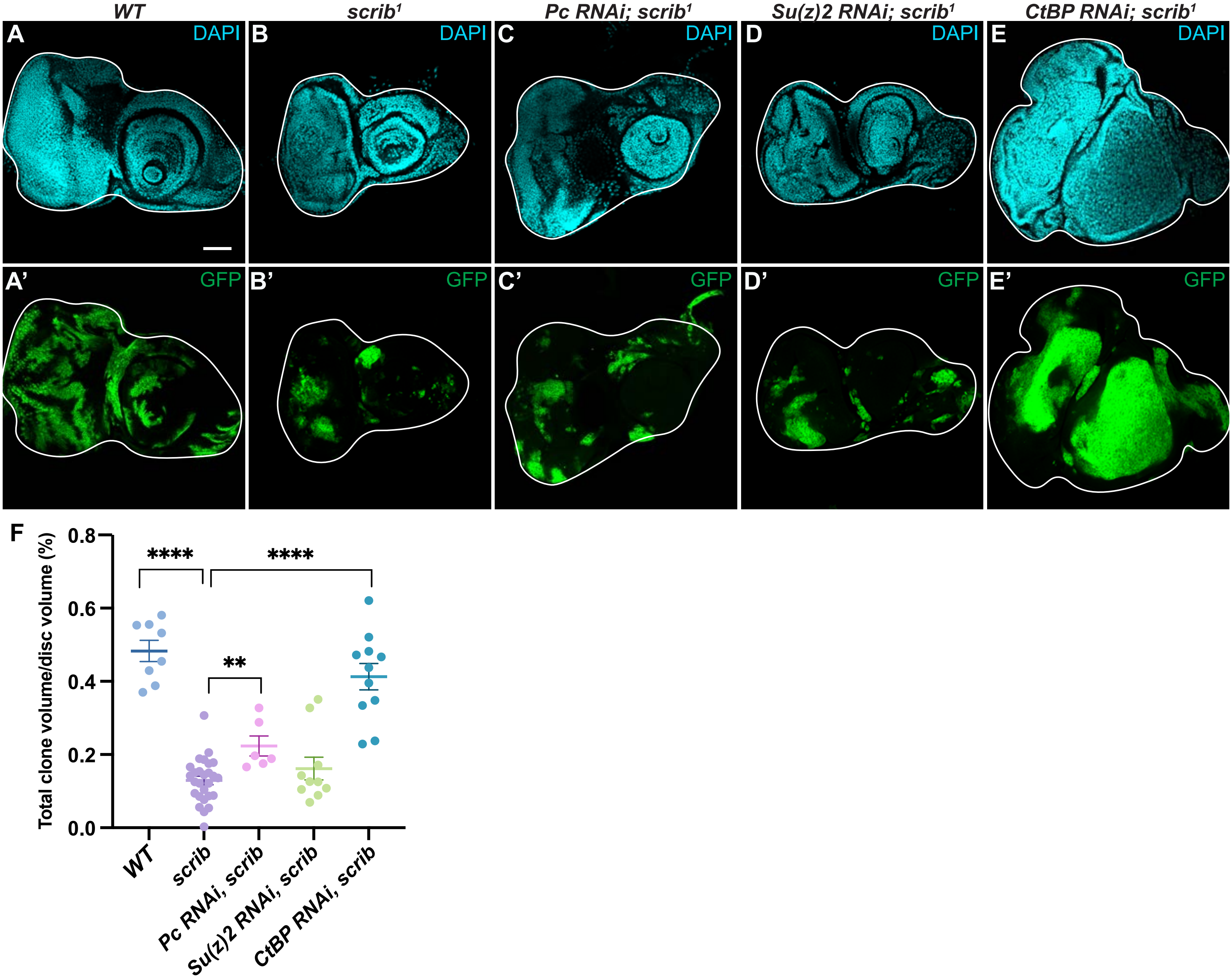

**Mitchell *et al.*, Fig. S4**

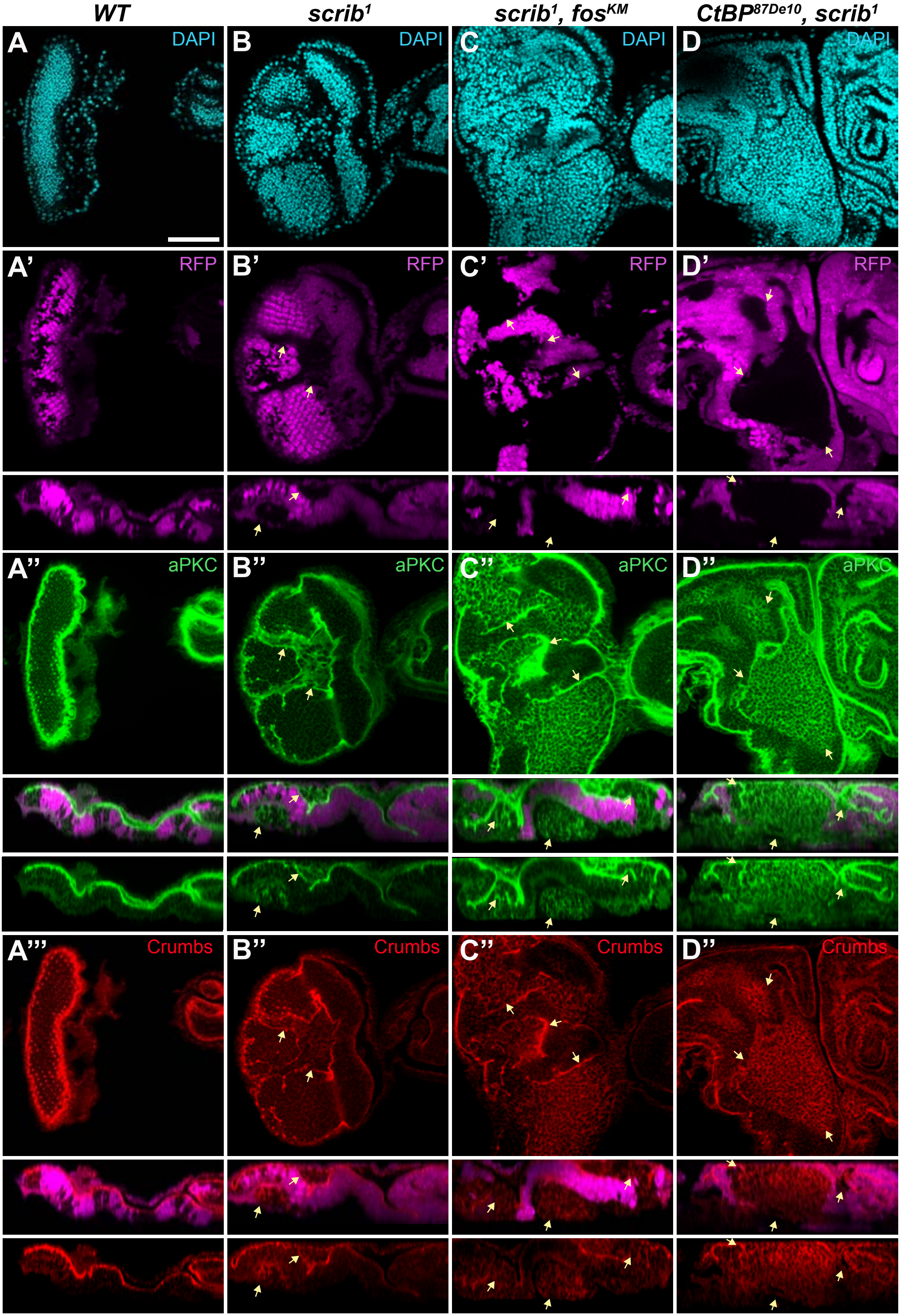

**Mitchell *et al.*, Fig. S5**

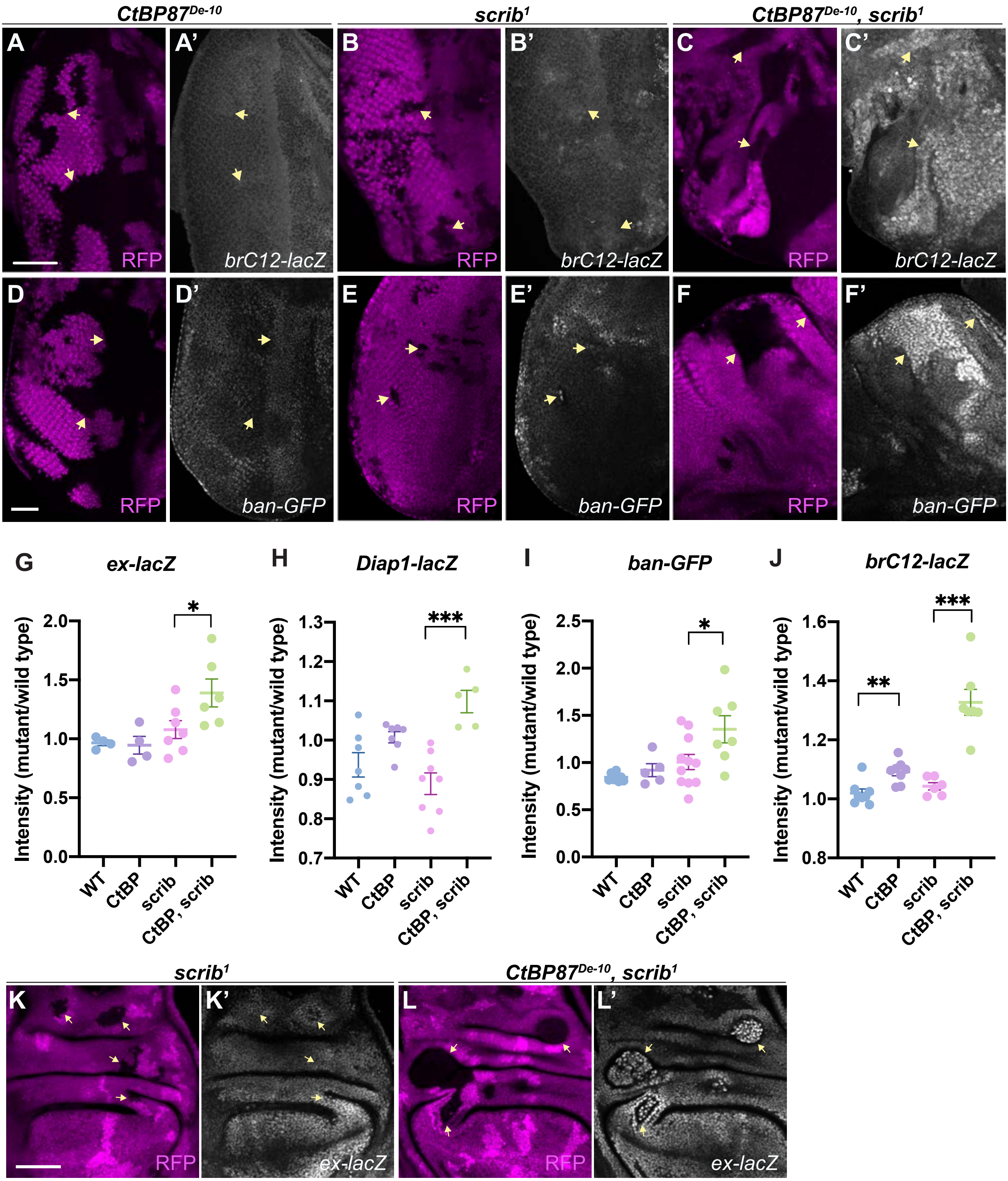

**Mitchell *et al.*, Fig. S6**

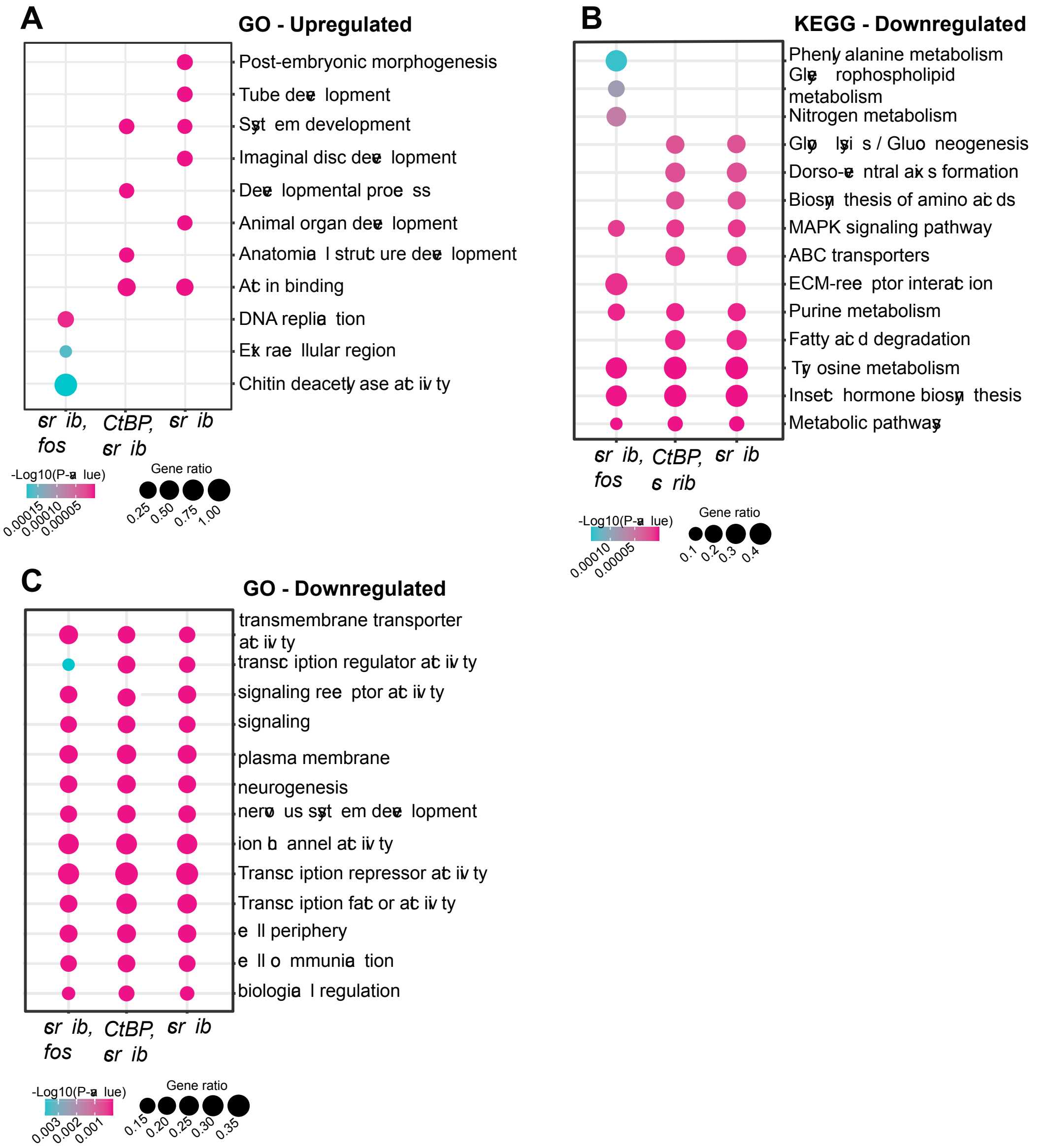

### Mitchell *et al.*, Fig. S7

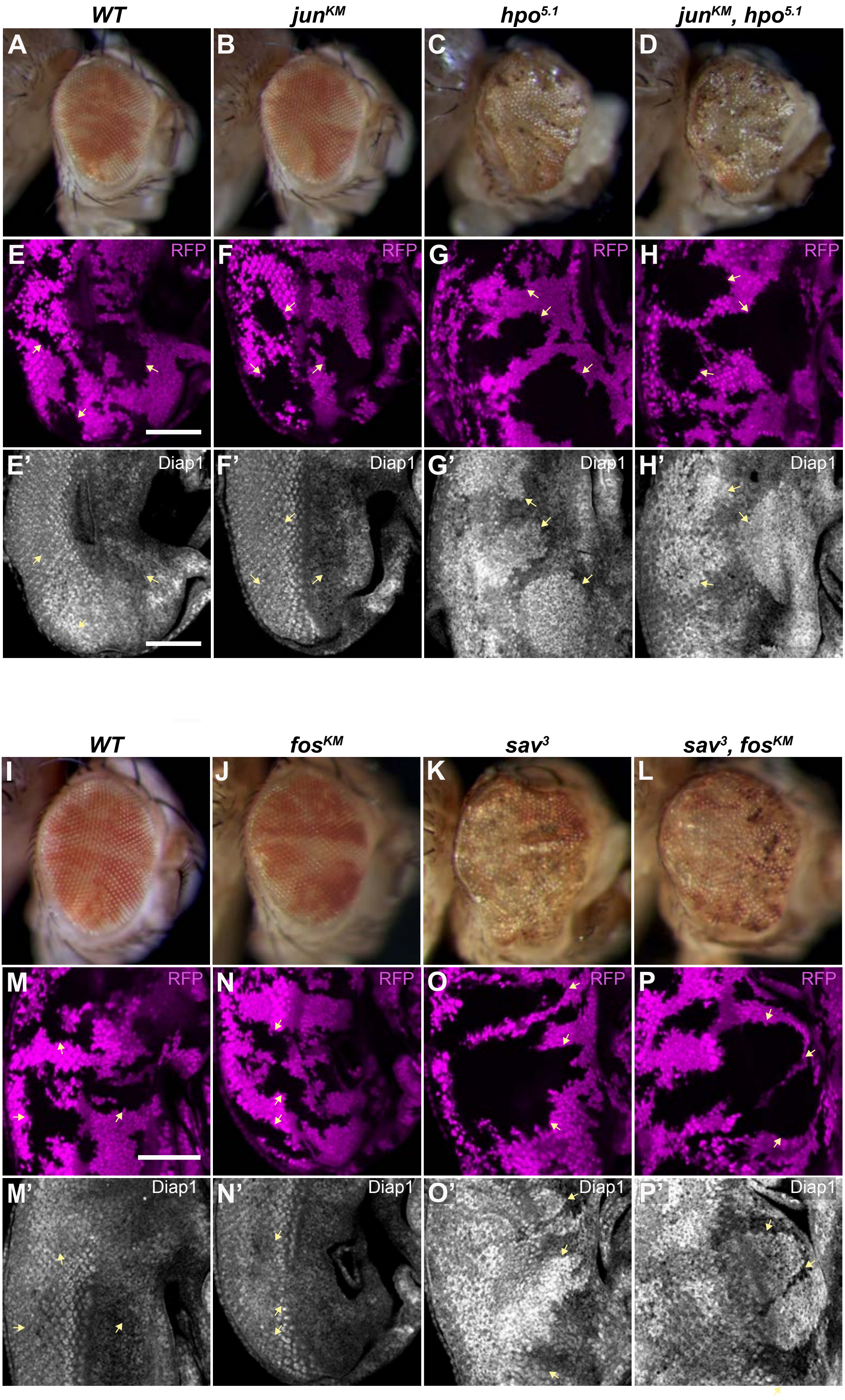

**Mitchell *et al*., Fig. S7**

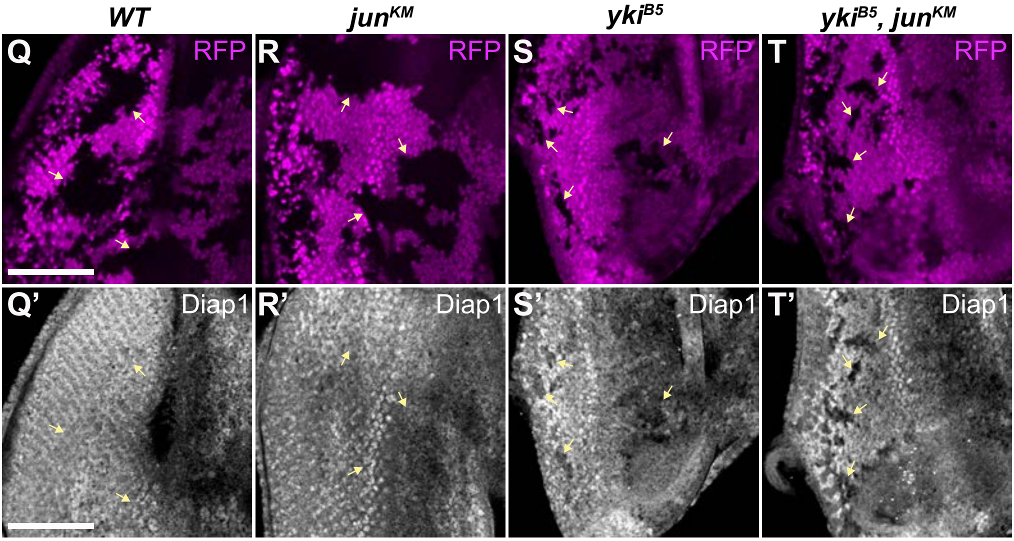
